# Direct observation of RAG recombinase recruitment to chromatin and the *IgH* locus in live pro-B cells

**DOI:** 10.1101/2020.09.07.286484

**Authors:** Geoffrey A Lovely, Fatima-Zohra Braikia, Amit Singh, David G. Schatz, Cornelis Murre, Zhe Liu, Ranjan Sen

## Abstract

The RAG1 and RAG2 proteins introduce double-strand DNA breaks at antigen-receptor loci in developing lymphocytes to initiate V(D)J recombination. How RAG proteins find the correct target locus in a vast excess of non-specific chromatin is not known. Here we measured dynamics of RAG1/RAG2 interactions with chromatin in living pro-B cells. We found that the majority of RAG1 or RAG1/RAG2 complex is in a fast 3D diffusive state, and the residual slow diffusive (bound) fraction was determined by a non-core portion of RAG1, and the PHD domain of RAG2. The RAG proteins exhibited distinct dynamics at the *IgH* locus. In particular, RAG2 increased the probability of RAG1 binding to *IgH*, a property that likely explains its non-catalytic role in V(D)J recombination. Our observations reveal how RAG finds its targets in developing B cells.

**One Sentence Summary:** Single-molecule imaging of the RAG recombinase reveals its search strategy for chromatin, H3K4me3 and antibody gene loci in living cells.

---

The immunoglobulin heavy chain gene (*IgH*) locus encodes the heavy chain of antibody molecules (*1*). Unlike most genes, functional *IgH* genes are assembled in somatic cells from widely dispersed variable (V_H_), diversity (D_H_) and joining (J_H_) gene segments by a process known as V(D)J recombination. Critical parts of the V(D)J recombination machinery are the lymphocyte-specific recombination activating gene products (RAG)1 and 2 (*2*, *3*). RAG1 and RAG2 together initiate recombination by recognizing recombination signal sequences (RSSs) associated with the gene segments and introducing double-strand DNA breaks (*2*, *3*). Thereafter, ubiquitously expressed proteins of the non-homologous end joining (NHEJ) pathway complete the reaction to produce functional genes (*2*, *3*). Being a DNA damaging agent, RAG1/RAG2 activity must be tightly regulated to minimize genomic instability. Indeed, off target effects of RAG1/RAG2 have been implicated in chromosomal translocations associated with human lymphomas and leukemias (*4*–*6*). Chromatin immunoprecipitation studies show that RAG1/RAG2 are localized at restricted regions of antigen receptor genes that are referred to as recombination centers (RCs) (*7*). These regions are also enriched for trimethylated lysine 4 on histone H3 (H3K4me3) and RAG1/RAG2 may localize here via a H3K4me3-binding plant homeodomain (PHD) in RAG2 (*8*). Additionally, RAG1/RAG2, and RAG2 alone can be found elsewhere in the genome at sites of high H3K4me3 enrichment (*8*, *9*). However, dynamic mechanisms by which RAG1/RAG2 find their genomic targets remain unclear.

To visualize RAG1 dynamics in live pro-B cells we fused a Halo tag at its N-terminus and expressed the fusion protein in an Abelson virus transformed pro-B cell line via retroviral transduction (Fig. 1A and fig. S1B) (*10*–*13*). Halo-RAG1 was expressed at levels comparable to endogenous RAG1 in the thymus (Fig. 1B, fig. S2, A and B), associated with endogenous RAG2 (fig. S3), and was functional in V(D)J recombination (fig. S4, A to C) (*14*). Having determined that the Halo tag did not affect RAG1 function, we initiated live cell imaging assays. To determine RAG1 diffusivity, we conducted two dimensional (2D) tracking experiments in 6312 pro-B cells with a 10ms integration time in order to simultaneously track RAG1 that is bound and diffusing (Fig. 1C) (*10*, *11*). We measured Halo-RAG1 trajectories in ≥ 9 cells in three independent experiments. As a control we measured histone H2B-Halo trajectories in 6312 cells (fig. S1H) (*10*, *11*). From these trajectories we calculated diffusion coefficients for Halo-RAG1 and H2B-Halo molecules. We found the RAG1 diffusion coefficient cumulative distribution was dominated by faster diffusive states compared to H2B (Fig. 1D). The range of diffusion coefficients (D) for RAG1 presumably reflects RAG1 that is diffusing and chromatin bound. To determine the fraction of RAG1 present in a bound state we compared its diffusivity to that of H2B. We fit the H2B diffusion coefficient distributions to a two gaussian mixture model (Fig. 1E), and determined a threshold slow diffusion coefficient of D = 0.05 ± 0.02μm^2^/s as a measure of the bound state (Fig. 1F).

**Fig. 1.**
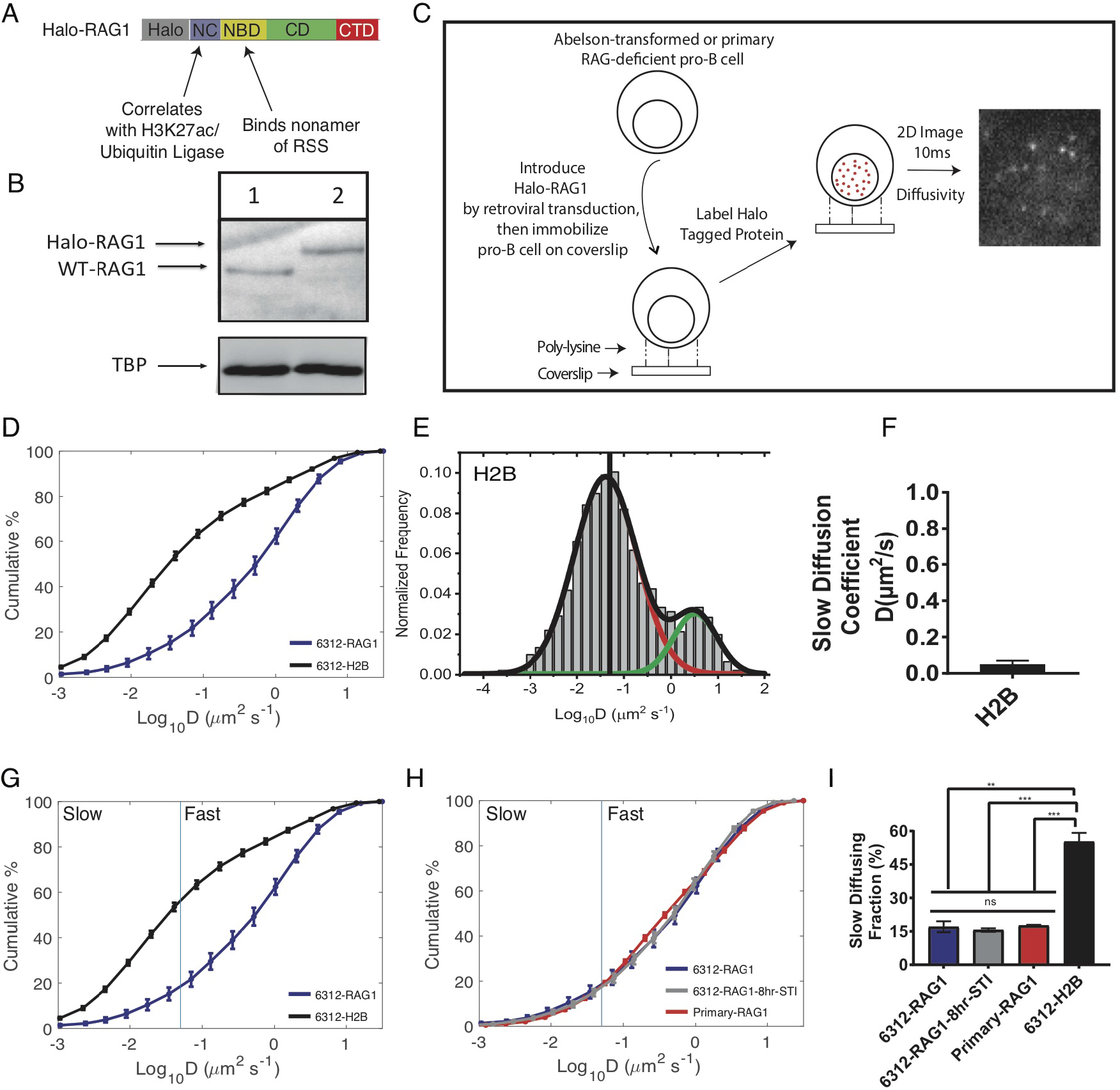
Single molecule imaging of RAG1 dynamics in live pro-B cells. (**A**) Schematic of Halo-RAG1 non-core domain (NC), nonamer binding domain (NBD), central domain (CD) and c-terminal domain (CTD). Domains that perform ubiquitylation, correlate with H3K27ac, and interact RSSs are highlighted. (**B**) Expression of Halo-RAG1 in 6312 pro-B cells (lane 2), and WT-RAG1 expression in mouse thymus (lane 1). Tata binding protein (TBP) was used as a loading control. (**C**) Live pro-B cell single-molecule imaging workflow to detect Halo-RAG1 freely diffusing and bound. (**D**) Cumulative distribution function (CDFs) of the diffusion coefficients for 6312-Halo-RAG1 (blue, n_cells_ = 35, n = 5695 trajectories) and 6312-H2B-Halo (black, n_cells_ = 18, n = 4806 trajectories). (**E**) Diffusion coefficient distribution with Gaussian fits for 6312-H2B-Halo (slow diffusion: red curve, and fast diffusion: green curve). (**F**) Average slow diffusion coefficient for 6312-H2B-Halo (black) from three independent experiments. The error bar represents the SEM. (**G**) CDFs of 6312-Halo-RAG1, and 6312-H2B-Halo. The fraction of the CDF to the left of the blue vertical line is slow diffusing. The blue vertical line is the slow diffusion coefficient for histone H2B. (**H**) CDFs of Halo-RAG1 diffusivity in 6312 cells (blue), 6312-cells treated with STI-571 for 8h (gray, n_cells_ = 25, n = 6945 trajectories), and in primary RAG2-deficient pro-B cells (red, n_cells_ = 24, n = 4523 trajectories). (**I**) Mean slow diffusing fractions for 6312-Halo-RAG1, 6312-Halo-RAG1-8hr STI, and RAG2-deficient primary pro-B-Halo-RAG1 and 6312-H2B-Halo. (All error bars represent SEM, **p < 0.01 ***p < 0.001.)

We then defined the fraction of RAG1 with D ≤ 0.05μm^2^/s as bound or slow diffusing and the fraction with D > 0.05μm^2^/s as 3D or fast diffusing (Fig. 1G). This approach was used previously to define the fraction of Cas9 in a bound or 3D diffusive state (*11*). We found 17.0 ± 2.5% of RAG1 to be in a slow diffusing state (Fig. 1, G to I) contrasting with the majority of H2B (55.3 ± 3.9%) in a slow diffusing state (Fig. 1, G and I) (*10*, *11*). The proportion of RAG1 in the slow and fast diffusive states were comparable in non-cycling 6312 (Fig. 1, H and I, fig. S5, A and B), and RAG2-deficient primary pro-B cells (Fig. 1, H and I, fig. S5, C and D) (*15*). We conclude that the bulk of nuclear RAG1 is in a 3D diffusive state in pro-B cells.

RAG1 contains a non-core (NC) domain that has been shown to be a stronger recruiter of RAG1 to chromatin genome-wide than the nonamer binding domain (NBD), which interacts with RSSs (*16*). Presence of the NC domain correlates with localization of RAG1 to histone H3K27ac modification, and also harbors ubiquitin ligase activity (*16*, *17*). To determine the extent to which the RAG1 NC and NBD domains contribute to RAG1 diffusivity, we measured diffusion coefficient distributions of RAG1 mutants, 6312-Halo-RAG1(ΔNBD) and 6312-Halo-RAG1(ΔNC) (Fig. 2A, fig. S1, C and D, fig. S2A) (*4*, *18*). We found that the ΔNBD did not change the slow diffusing fraction compared to RAG1, however deleting the NC domain resulted in a 3-fold reduction in this fraction (5.6 ± 0.6%) (Fig. 2, B and C). It is possible that the reduction in the bound fraction of RAG1(ΔNC) resulted from significantly greater expression of this protein compared to RAG1 (fig. S2A). By fitting the diffusion coefficient distribution with a two gaussian mixture model (Fig. 2D), we found no significant difference in the slow diffusion coefficients for RAG1 (0.16 ± 0.06μm^2^/s) and RAG1(ΔNBD)(0.26 ± 0.11μm^2^/s), however this was increased 4-fold for RAG1(ΔNC)(0.74 ± 0.19 μm^2^/s) (Fig. 2E). We conclude that the NC rather than the NBD domain slows down the 3D diffusion of RAG1, presumably by interacting with chromatin. This is consistent with the observation that the RAG1 NC domain, and not the NBD is required for RAG1 chromatin interactions measured by ChIP (*6*).

**Fig. 2.**
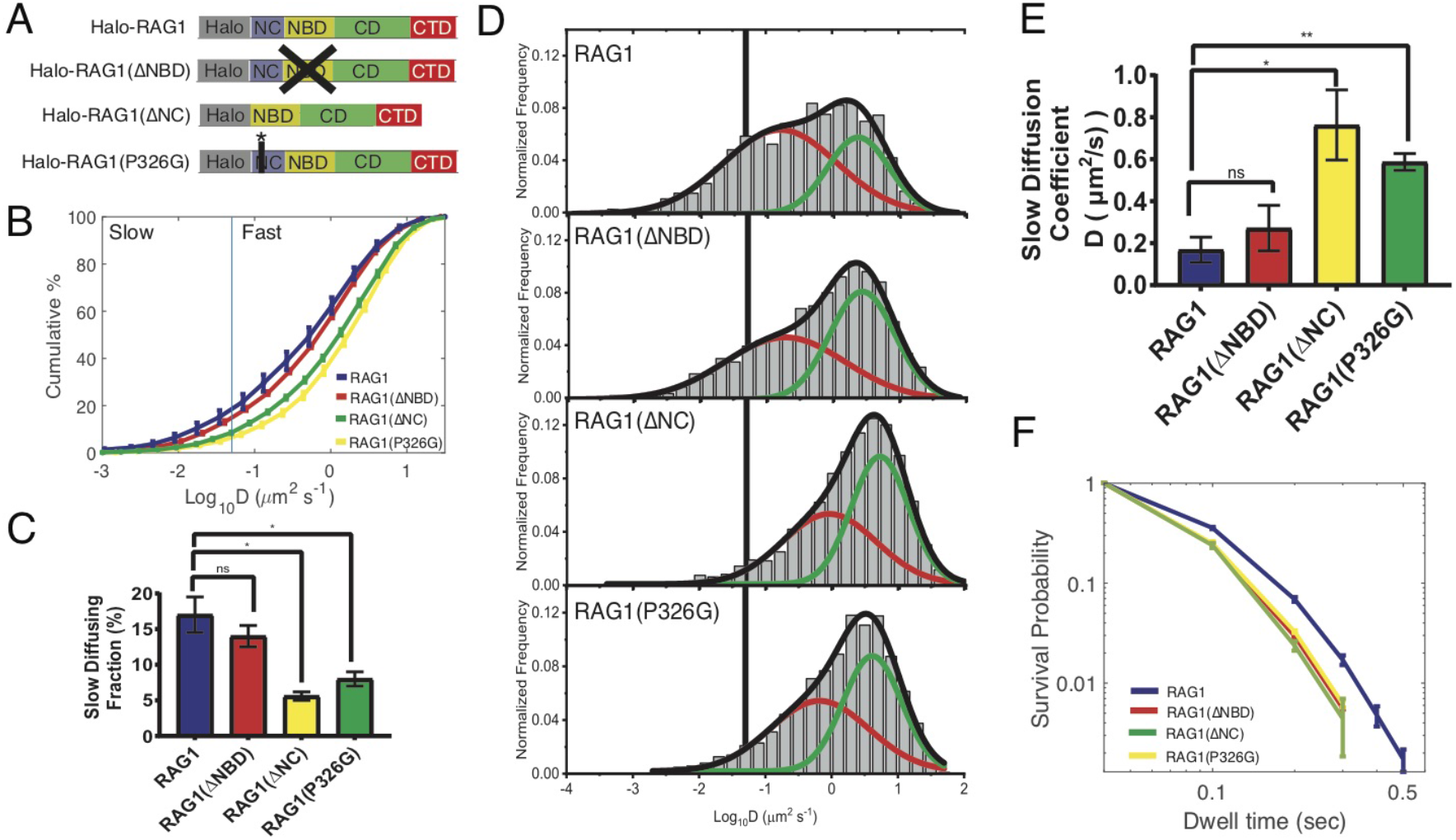
Contribution of RAG1 non-core domain to diffusion dynamics in pro-B cells. (**A**) Schematics of Halo-RAG1, Halo-RAG1with mutations in residues R391A, R393A, R402A (Halo-RAG1(ΔNBD)), Halo-RAG1 truncation mutation containing residues 384-1040 (Halo-RAG1(ΔNC)), and Halo-RAG1 with a mutation in residue P326G (Halo-RAG1(P326G)). (**B**) CDFs of diffusion coefficient distributions for Halo-RAG1(blue, n_cells_ = 35, n = 5695 trajectories), Halo-RAG1(ΔNBD) (red, n_cells_ = 22, n = 5506 trajectories), Halo-RAG1(ΔNC) (yellow, n_cells_ = 23, n = 4128 trajectories) and Halo-RAG1(P326G) (green, n_cells_ = 15, n = 6055 trajectories). The fraction of the CDF to the left of the blue vertical line is slow diffusing. The blue vertical line is the slow diffusion coefficient for histone H2B. (**C**) Mean slow diffusing fraction for Halo-RAG1, Halo-RAG1(ΔNBD, Halo-RAG1(ΔNC) and Halo-RAG1(P326G). (**D**) Representative diffusion coefficient distribution and gaussian fits for Halo-RAG1, Halo-RAG1(ΔNBD), Halo-RAG1(ΔNC) and Halo-RAG1(P326G). The fraction of the histograms to the left of the black vertical line are slow diffusing based on the histone H2B slow diffusion coefficient cutoff. (**E**) Mean slow diffusion coefficient for Halo-RAG1, Halo-RAG1(ΔNBD), Halo-RAG1(ΔNC) and Halo-RAG1(P326G). (**F**) Dwell-time distributions for Halo-RAG1 (blue, n_cells_ = 9, n = 4083 trajectories), Halo-RAG1(ΔNBD) (red, n_cells_ = 15, n = 4838), Halo-RAG1(ΔNC) (yellow, n_cells_ = 23, n = 3618 trajectories) and Halo-RAG1(P326G) (green, n_cells_ = 17, n = 4519).(All error bars represent SEM, *p < 0.05, **p < 0.01.)

To more precisely probe the effect of the ubiquitin ligase domain on RAG1 dynamics we measured diffusion parameters of a RAG1 derivative with the P326G point mutation in this domain (Fig. 2A, fig S1E, fig. S2A) (*18*, *19*). This mutation has been previously shown to inactivate E3 ligase activity *in vitro* (*19*). We found that the slow diffusing fraction was reduced (Fig. 2, B and C), and the corresponding diffusion coefficient increased by this mutation (Fig. 2, D and E) in a manner comparable to RAG1(ΔNC). Thus, a large part of RAG1 chromatin interactions appears to require ubiquitin ligase activity.

To determine if the different domains of RAG1 contributed to the duration of protein binding to chromatin, we measured the dwell-time of RAG1 and RAG1 mutants using an approach established previously (*11*, *12*, *20*). Briefly, we imaged continuously with a 20ms integration time and changed the maximum expected diffusion coefficient during the analysis to 0.05μm^2^/s (*11*, *12*, *20*). We found that the length and survival probability of RAG1 dwell-times was reduced upon mutation of the NBD domain, NC domain or residue P326G (Fig. 2F), indicating that the RAG1 NBD, NC and ubiquitin ligase domains play a role in regulating the time RAG1 spends bound to chromatin genome-wide.

Next, we investigated the effect of RAG2 on RAG1 dynamics by introducing RAG2 into the 6312-Halo-RAG1 pro-B cell line by lentiviral transduction (fig. S6, Fig. 3A). The introduced RAG2 formed a complex with Halo-RAG1 (fig. S3) that permitted DJ recombination on endogenous *IgH* alleles (Fig. 3B, lanes 4-6). From the diffusion coefficient distribution of 6312-Halo-RAG1/RAG2, we found 14.6 ± 0.6% of the complex to be in the slow diffusing fraction (Fig. 3, C and D). Thus, the presence of RAG2 did not significantly change the proportion of RAG1 in the slow and fast diffusing fractions. However, point mutation of the PHD domain of RAG2(W453A)(Fig. 3A), that binds poorly to H3K4me3 (*8*), and abrogated DJ recombination (Fig. 3B, lanes 7-9), significantly reduced the slow diffusing fraction (10.0 ± 0.5%) (Fig. 3, C and D). Applying the two Gaussian fit strategy to diffusion coefficient distributions we found no significant difference between the slow diffusion coefficients of RAG1/RAG2, and RAG1/RAG2(W453A) (Fig. 3E, fig. S7). These complexes also did not show any significant difference in their dwell-time distributions (Fig. 3F). We conclude that the RAG2 PHD domain regulates the fraction of RAG1/RAG2 complex bound to chromatin, but not the time spent bound. Our interpretation of these observations is that interaction of the RAG1/RAG2 complex with bulk chromatin is skewed towards recognition of H3K4me3 via the RAG2 PHD domain. Reduction of the bound fraction of RAG1/RAG2(W453A) below that of RAG1 alone suggests that RAG2 alters RAG1’s interaction in ways currently not understood, resulting in coincidentally similar values for the bound fractions of RAG1 alone or the RAG1/RAG2 complex. We observed no significant difference in RAG1/RAG2 dynamics in non-cycling 6312 cells (*21*) (fig. S8, A and B), nor in RAG1-deficient primary pro-B cells (fig. S8, A and B, fig. S9, A to C). Together these data show that RAG1/RAG2 primarily uses 3D diffusion to scan the genome and bind to sites with H3K4me3 in pro-B cells.

**Fig. 3.**
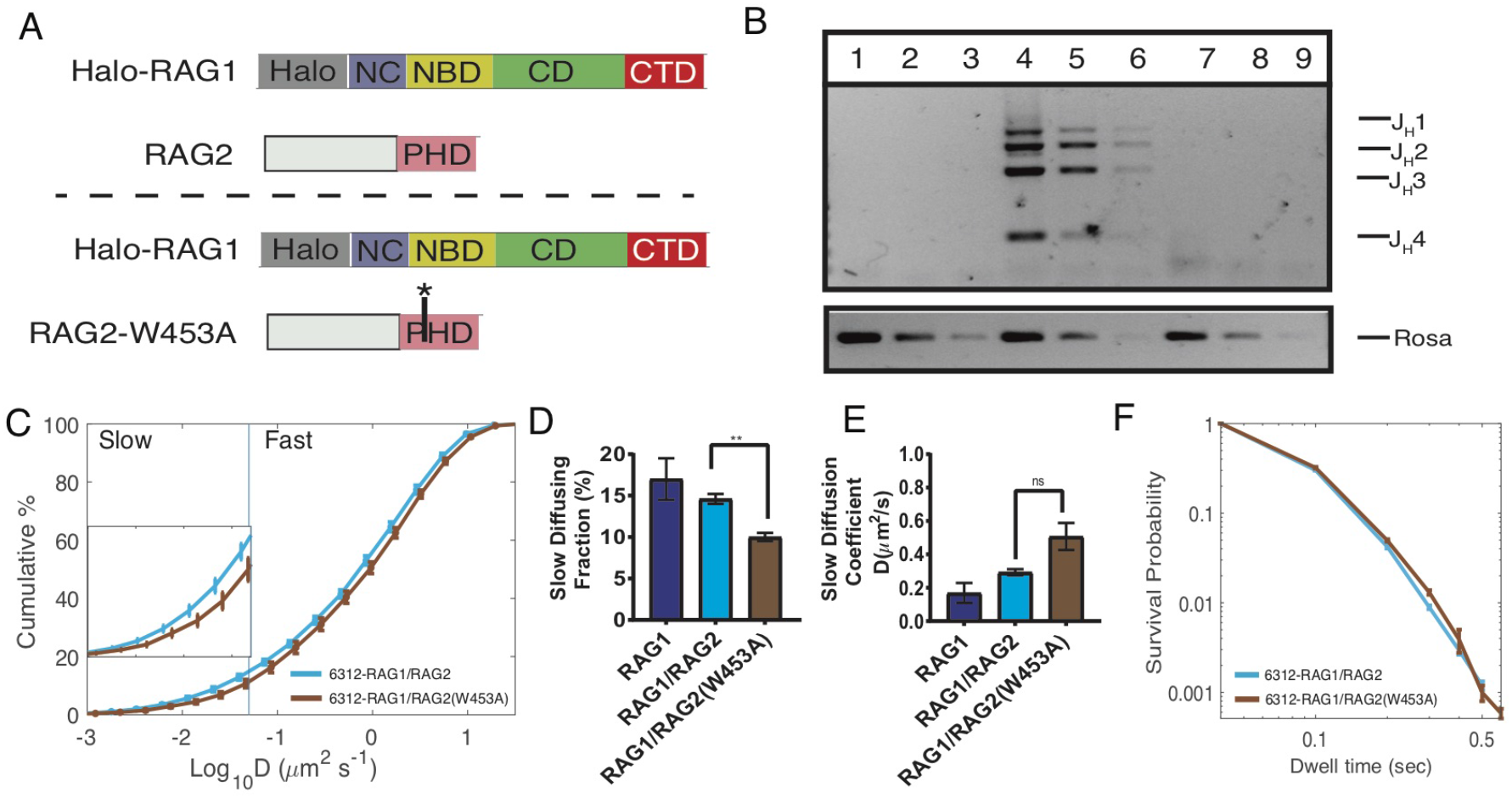
Effect of RAG2 on RAG1 dynamics in pro-B cells. (**A**) Schematics of Halo-RAG1 in the presence of unlabeled RAG2, and RAG2 with a point mutation in residue W453A. (**B**) DJ recombination assay on 6312-Halo-RAG1 (lanes: 1-3), 6312-Halo-RAG1/RAG2 (lanes: 4-6), 6312-Halo-RAG1/RAG2(W453A) (lanes:7-9). (**C**) CDFs of Halo-RAG1/RAG2 (cyan, n_cells_ = 16, n = 6880 trajectories) and Halo-RAG1/RAG2-W453A (brown, n_cells_ = 23, n = 5923 trajectories). The fraction of the CDF to the left of the blue vertical line is slow diffusing. The blue vertical line is the slow diffusion coefficient for histone H2B. (**D**) Mean slow diffusing fraction of Halo-RAG1, Halo-RAG1/RAG2, and Halo-RAG1/RAG2-W453A. (**E**) Mean slow diffusion coefficient of Halo-RAG1/RAG2, and Halo-RAG1/RAG2-W453A. (**F**) Dwell-time distributions for Halo-RAG1/RAG2 (cyan, n_cells_ = 16, n = 8590 trajectories) and Halo-RAG1/RAG2-W453A (brown, n_cells_ = 25, n = 8581 trajectories). (All error bars represent SEM, **p < 0.01.).

Lastly, we sought to distinguish the dynamics of bulk nuclear RAG1/RAG2 from RAG1/RAG2 molecules at a target antigen receptor locus (Fig. 4A). For this we used RAG2-deficient KM-Tet-O pro-B cell line that harbors 240 Tet operator sites located 3.1kb from the recombination center on *IgH* alleles (*22*). We first introduced Tet-R-GFP (*23*) to mark *IgH* alleles, then imaged with a 10ms integration time to measure their diffusivity compared to histone H2B. We found that the diffusion coefficient of the *IgH* locus (0.14 ± 0.01μm^2^/s) (fig. S10, A to D) was greater than that of histone H2B (0.05 ± 0.02μm^2^/s) (Fig. 1, E and F, fig. S10, B to D). We ruled out localization uncertainty caused by the array of bound Tet-R-GFP as a factor contributing to the increased diffusivity at the *IgH* locus relative to H2B (see methods). Since the motion of both H2B (in the form of nucleosomes) and Tet-R-GFP represent chromatin movements, we infer that the *IgH* locus in pro-B cells is more dynamic than bulk nuclear chromatin. Previous studies have shown that measurements of individual *IgH* alleles maybe confounded by nuclear and cellular rotations (*23*). Because these parameters acted equally on H2B-containing chromatin and *IgH* alleles, we infer that the faster diffusivity of *IgH* reflects the unique chromatin structure of an active locus, compared to bulk chromatin. This increase in motility may contribute to the recruitment of RAG1/RAG2 to the *IgH* locus in pro-B cells.

**Fig. 4.**
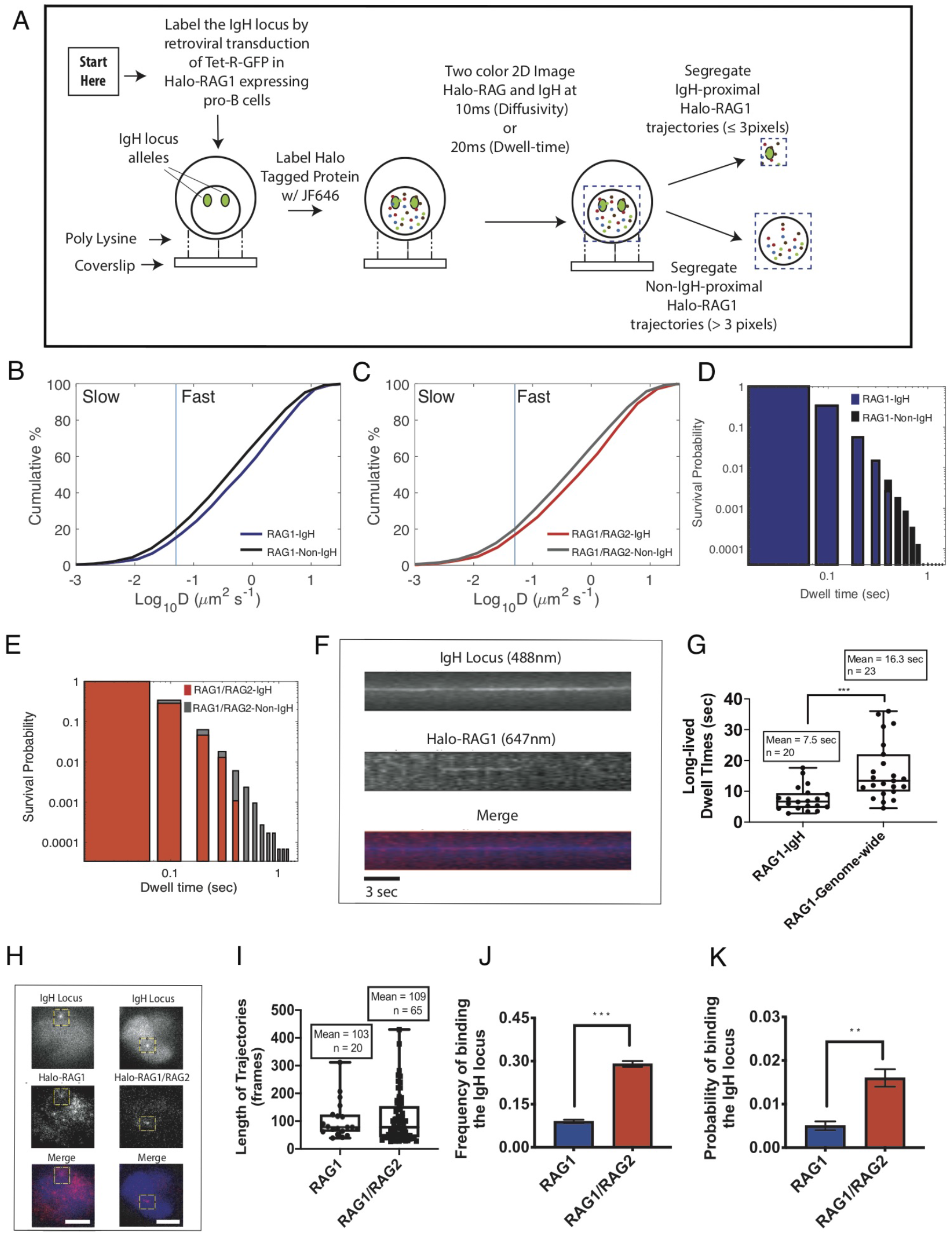
RAG1 and RAG1/RAG2 dynamics at the *IgH* locus in live pro-B cells. (**A**) Schematic of workflow to performing two color imaging of Halo-RAG1 or Halo-RAG1/RAG2 and the *IgH* locus in live pro-B cells. (Note: Although both *IgH* alleles are shown in the schematic, the majority of cells analyzed only had one *IgH* allele in focus). (**B**) CDFs of diffusion coefficients for *IgH*-proximal RAG1 trajectories (blue) and non-*IgH*-proximal RAG1 trajectories (black) (n_cells_ = 270, n_RAG1-IgH_ = 1034 trajectories, n_RAG1-Non-IgH_ = 13,670 trajectories). (**C**) CDFs of diffusion coefficients for *IgH*-proximal RAG1/RAG2 trajectories (red) and non-*IgH*-proximal RAG1/RAG2 trajectories (gray) (n_cells_ = 237, n_RAG1/RAG2-IgH_ = 974 trajectories, n_RAG1/RAG2-Non-IgH_ = 16,497 trajectories). The fraction of the CDF to the left of the blue vertical line is slow diffusing. The blue vertical line is the slow diffusion coefficient for histone H2B. (**D**) Dwell-time distributions of *IgH*-proximal RAG1 trajectories (blue bars) and non-*IgH*-proximal RAG1 trajectories (black bars) (n_cells_ = 117, n_RAG1-IgH_ = 1153 trajectories, n_RAG1-Non-IgH_ = 24,584 trajectories), and (**E**) *IgH*-proximal RAG1/RAG2 trajectories (red bars) and non-*IgH*-proximal RAG1/RAG2 trajectories (gray bars) (n_cells_ = 154, n_RAG1/RAG2-IgH_ = 927 trajectories, n_RAG1/RAG2-Non-IgH_ = 29,418 trajectories). (**F**) Vertical slice of long-lived Halo-RAG1 (blue), colocalized with the *IgH* locus (red) that is assembled horizontally frame by frame through time into a kymograph. (**G**) The long lived mean dwell-time for Halo-RAG1 is determined by imaging with a 20ms integration time, followed by a 300ms darktime. The 300ms darktime increases the survival of the fluorescent probe yielding a long-lived measure of the dwell-time. A long-lived dwell-time was determined for RAG1 at the *IgH* locus (n_cells_ = 15, n = 20 interactions), and genome-wide (n_cells_ = 6, n = 23 interactions) (***p < 0.001). (**H**) Colocalization of bound Halo-RAG1 or Halo-RAG1/RAG2 at the *IgH* locus after 10ms two-color imaging (scale bar = 5*μ*m). (**I**) Length of RAG1 (n_cells_ = 195, n_IgH alleles_ = 209, n = 20 interactions) and RAG1/RAG2 (n_cells_ = 195, n_IgH_ alleles = 211, n = 65 interactions) trajectories co-localized with the *IgH* locus in frames used that were used to compute the frequency, and probability of binding to the *IgH* locus. (**J**) The frequency of RAG1 or RAG1/RAG2 binding to the *IgH* locus is computed by dividing the number of co-localization events, by the number of *IgH* alleles assayed (3 independent experiments, n = 65 cells per/experiment, error bars represent SEM, and ***p < 0.001.). (**K**) The probability of RAG1 or RAG1/RAG2 binding to the *IgH* locus is computed by dividing the time RAG spends bound to the *IgH* locus by the total time the *IgH* locus is bound or unbound. The total time was set to 2,000 frames, because no RAG1 or RAG1/RAG2 binding to the *IgH* locus was detectable after this cutoff. Therefore the only parameter changing in the equation is the time the *IgH* locus is bound by RAG1 or RAG1/RAG2 (3 independent experiments, n = 65 cells per/experiment, error bars represent SEM, and **p < 0.01.).

Next, we introduced Halo-RAG1 into KM-Tet-O pro-B cells with or without RAG2 by retroviral transduction (Fig. 4A, fig. S11, A to C, fig. S12, A to C). After enriching Halo-RAG1 expressing cells by flow cytometry we expressed Tet-R-GFP to illuminate *IgH* alleles (Fig. 4A, fig. S13, A and B), and carried out two-color imaging to measure Halo-RAG1 or Halo-RAG1/RAG2 diffusivity and dwell-times (*12*).

As a first level of distinction we compared Halo-RAG1 trajectories within 3 pixels (480nm) of the *IgH* locus to the bulk distribution (Fig. 4A) (*12*). We found the fraction of slow diffusing *IgH*-proximal RAG1(15%) or RAG1/RAG2 (16%) trajectories was reduced relative to non-*IgH*-proximal RAG1 (19%) or RAG1/RAG2 (20%) trajectories (Fig. 4, B and C). We also found the slow diffusion coefficient was increased for *IgH*-proximal RAG1 (0.43μm^2^/s) or RAG1/RAG2 (0.44μm^2^/s) trajectories compared to non-*IgH*-proximal RAG1 (0.29μm^2^/s) or RAG1/RAG2 (0.27μm^2^/s) trajectories (fig. S14, A to D). This indicates that RAG1 or RAG1/RAG2 moves faster in the vicinity of the *IgH* locus in pro-B cells compared to its diffusivity genome-wide. Additionally, RAG1 or RAG1/RAG2 dwell-times were reduced at *IgH*-proximal compared to non-*IgH*-proximal trajectories (Fig. 4, D and E).

To more closely examine RAG1 dynamics at the *IgH* locus we used the reslice, merge and measure tools in ImageJ to only investigate Halo-RAG1 trajectories that colocalized with the *IgH* locus (Fig. 4F) (*12*). We determined a measure of the long-lived dwell-time of Halo-RAG1 at the *IgH* locus by imaging with a 20ms integration time, followed by a 300ms dark time (*11*, *12*). We found the long-lived mean dwell-time of Halo-RAG1 at the *IgH* locus is 7.5 secs (Fig. 4G). This is shorter than the 16.3 sec long-lived mean dwell-time for Halo-RAG1 determined genome-wide (Fig. 4G) (*10*).

Lastly, we used the reslice, merge and measure tools again, but on the 10ms two-color imaging data (Fig. 4H) to identify Halo-RAG1 or Halo-RAG1/RAG2 co-localized with the *IgH* locus. The 10ms integration time permitted visualization of shorter duration contacts of RAG with *IgH*. This approach allowed us to compute the frequency of RAG1 or RAG1/RAG2 binding to the *IgH* locus. For this we divided the number of colocalization events by the total number of *IgH* alleles assayed (Fig. 4I). We detected a 3-fold increased frequency of RAG1 binding to *IgH* in the presence of RAG2 (Fig. 4J). We also calculated a probability of RAG binding to the *IgH* locus as the ratio of time that *IgH* was bound by RAG1 or RAG1/RAG2 to the total recording time (Fig. 4I, fig. S15, A and B) (*24*). We observed a 3-fold increased probability of RAG1 binding to *IgH* in the presence of RAG2 (Fig. 4K). Since the mean length of trajectories, a measure of the off-rate, was the same for either RAG1 or RAG1/RAG2 (Fig. 4I), we infer that RAG2 increased the on-rate of RAG1 at the *IgH* locus. Furthermore, the low probability of RAG1/RAG2 binding to the *IgH* locus (0.016) (Fig. 4K) indicates that most of the time the *IgH* locus remains unoccupied. Together these data reveal a distinct target search strategy, dwell-time distribution, and probability of binding for RAG1 and RAG1/RAG2 at the *IgH* locus.

Our data provide a direct window into RAG’s target search strategy in live pro-B cells. RAG uses a 3D diffusion-dominated search strategy to interact with chromatin, H3K4me3 and the *IgH* locus. We found mutation of RAG non-core domains increased its diffusivity and reduced its dwell-time genome-wide, demonstrating a possible function of the ubiquitin ligase domain in this search strategy. We found that RAG2 increased the probability of RAG1 binding to the *IgH* locus by increasing RAG1’s on-rate, though the locus was mostly unoccupied. Once bound, RAG1 alone remained bound for roughly 7.5 sec and we inferred that the dwell-time of the RAG1/RAG2 complex is comparable. These observations explain in part the essential non-catalytic role of RAG2 in V(D)J recombination.

Chromatin loop extrusion has recently been shown to be the main process directing RAG-mediated V(D)J recombination (*25*, *26*). The first step of *IgH* gene assembly occurs in a 60kbp domain that includes DH gene segments, and the recombination center (*25*). Based on recently determined *in vitro* rates of chromatin loop extrusion of 0.5kbp/sec (*27*) the DH gene segments would take ~120 sec to pass by the recombination center. Since RAG spends 7.5 sec bound to the *IgH* locus, within the time it takes to extrude the 60kbp region RAG could bind at most ~16 times, assuming a probability of binding of 1. However, once we use the RAG1/RAG2 probability of binding the *IgH* locus (0.016), the number of binding events reduces significantly to ~0.27 times. Therefore, the 60kbp domain at the *IgH* locus can undergo chromatin loop extrusion 4 times before a RAG1/RAG2 binding event. This suggests a model for V(D)J recombination in which the *IgH* locus recombination center is infrequently bound by RAG while it undergoes chromatin loop extrusion. Mechanisms by which chromatin loop extrusion dynamics and direction are coordinated with sporadic presence of RAG proteins on *IgH* alleles to ensure fidelity of DH recombination await further studies. One possibility is that either RAG dynamics or loop extrusion dynamics (or both) are altered on alleles that simultaneously bind RAG1/RAG2 and cohesin.

## Acknowledgements

Thank you to the members of the Sen, Liu, Murre, and Schatz labs for helpful discussions. We would also like to thank Justin Bois for help with analysis, and Luke Lavis for generously providing dyes.

## Authors contributions

G.A.L., Z.L. and R.S. conceived and designed study. G.A.L. performed most of the experiments, F.Z.B performed DJ recombination assays, A.S. performed cell cycle distribution assays. C.M. and D.G.S provided essential reagents. G.A.L, F.Z.B, A.S and R.S. analyzed the data. G.A.L., Z.L. and R.S. interpreted the data and results and wrote the manuscript. All authors provided input and approved the final manuscript.

## Data and materials availability

All data needed to evaluate the conclusions in the paper are present in the paper and/or the Supplementary Materials. Additional data, and code related to this paper may be requested from the authors.

