## Supplementary Information for "Direct observation of RAG recombinase recruitment to chromatin and the *IgH* locus in live pro-B cells"

**This PDF file includes:**

Materials and Methods
Figs. S1 to S15
Table S1
Movie Legends S1 to S10

**Other Supplementary Materials for this manuscript include the following:**

Movies S1 to S10

**Materials and Methods**

Mice, Cell lines and Cell Culture.

6312-RAG2$^{-/-}$, DGS-RAG1$^{-/-}$ (a gift from David Schatz), KM-Tet-O-RAG2$^{-/-}$(a gift from Cornelis Murre) pro-B cell lines were cultured in RPMI-1640, 10% (6312 and DGS) or 20% (KM-Tet-O) FBS, 2mM L-glutamine, 100$\mu$g/mL penicillin, 100$\mu$g/mL streptomycin, 0.7X NEAA, 10mM HEPES (pH: 7.4) and 50$\mu$M 2-mercaptoethanol. RAG1 and RAG2-deficient mice were utilized to isolate primary pro-B cells via CD19 positive selection. The primary pro-B cells were cultured in Opti-MEM, 10% FBS, 2mM L-glutamine, 100$\mu$g/mL penicillin, 100$\mu$g/mL streptomycin, 0.7X NEAA, 10mM HEPES (pH: 7.4), 50$\mu$M 2-mercaptoethanol and 2ng/mL IL-7.

Isolating and expanding RAG-deficient primary pro-B cells from bone marrow

We isolate the tibia, and femurs from RAG-deficient mice, and extract the bone marrow. We then lyse the red blood cells, and perform CD19 positive selection on magnetic beads to isolate the primary pro-B cells. The isolated primary pro-B cells are then cultured in Opti-MEM, 10% FBS, 50$\mu$M 2-mercaptoethanol, 2mM L-glutamine, 100$\mu$g/mL penicillin, 100$\mu$g/mL streptomycin, 0.7X NEAA, and 2ng/mL IL-7, at a minimal density of 0.5 x $10^6$ cells/mL. The cells are cultured for a maximum of 7 days, and retroviral or lentiviral infections were performed on the 4th day of culture.

Retrovirus or Lentivirus Vectors, Assembly and Infections.

The Moloney murine leukemia retroviral vector system pOOC was a kind gift from Alejandro Balazs. All cloning was performed using Gibson assembly (NEB). To study RAG1 chromatin targeting we fused a Halo-tag to the N-terminus of various RAG1 mutants. The cloned RAG1 mutant retroviral vectors are pOOC-CMV-Halo-RAG1, pOOC-CMV-Halo-RAG1(ΔNBD)(R391A, R393A, R402A), pOOC-CMV-Halo-RAG1(ΔNC)(residues 384-1040) and pOOC-CMV-Halo-RAG1(P326G). To investigate unlabeled RAG2 we created pHIV-RAG2-IRES-Puro and pHIV-RAG2(W453A)-IRES-Puro. To label the *IgH* locus Tet-O array we obtained a LMP-Tet-R-EGFP retroviral vector (a kind gift from Cornelis Murre). Lentivirus was assembled by adding 45$\mu$g pHIV-viral genome, 33$\mu$g packaging vector, and 22$\mu$g VSVG envelope vector into 4mL of 0.22$\mu$m filtered DMEM (media only). Then the contents is mixed with 100$\mu$L of the transfection reagent BioT (Bioland), vortexed, and incubated at 25°C for 5 min. Then the 4mL of DMEM, BioT, viral genome/packaging/envelope vector mix is distributed, 1 mL each into 4 x 10cm dishes of 80% confluent 293T cells. The transfection proceeds for 2 days. To assemble retrovirus 60$\mu$g of viral genome vector, and 5$\mu$g VSVG was added to 4mL of filtered DMEM, and 100$\mu$L of BioT. Then the 4mL of DMEM, BioT, viral genome/envelope vector mix is distributed, 1 mL each into 4 x 10cm dishes of PLAT-E cells (CellBioLabs) that are 80% confluent. The transfection proceeds for 2 days. Then the lentiviral or retroviral supernatant is collected, 0.45$\mu$m filtered, and concentrated using ultracentrifugation (SW-32-Ti, 16,500 rpm, 1hr 30min). We remove the supernatant, then resuspend the pellet in 0.5 mL of RPMI complete media containing 10$\mu$g/mL polybrene. If infecting Abelson-transformed pro-B cells with retrovirus we add 500,000 cells to the 0.5mL of resuspended virus in a 24 well plate. After two days we either perform FACs or use antibiotic resistance. If infecting with lentivirus we spin infect the Abelson-transformed pro-B cells at 2,000 rpm for 40min @ 37°C. If we are infecting primary pro-B cells with retrovirus, on the 4th day of culture we resuspend the viral pellet in 0.5mL of Opti-MEM, then we add the 0.5mL to 4mL of Opti-MEM containing 2,000,000 cells at a density 500,000/mL. After two days we perform FACs.

Flow Cytometry.

Infected Abelson-transformed pro-B cell lines and primary pro-B cells were sorted using a BD FACS Aria Fusion. Halo-tag expressing pro-B cell line or primary cells were dyed using JF549 ligand (Gift from Luke Lavis) *(28)*.

DJ Recombination Detection Assays**.**

Genomic DNA was extracted from 6312, DGS, or KM-Tet-O pro-B cell lines harboring Halo-RAG1 with or without unlabeled RAG2 or RAG2(W453A) to confirm the halo-tag does not ablate V(D)J recombination. Then 60ng, 12ng or 2.4ng of genomic DNA was used per reaction, and amplified using a hotstar high fidelity polymerase (Qiagen). The PCR program used was 95°C: 15 min, 94°C: 45 sec, 60°C: 1 min, 72°C: 2 min, repeat (33 cycles), 72°C: 10 min. The *Rosa* locus was used as the loading control.

Western Blots.

To perform western blots on Abelson transformed pro-B cell lines, primary pro-B cells, and thymic cells, 20 X $10^6$ cells were lysed with 100$\mu$L of pro-B lysis buffer (20mM Tris pH 7.5, 20mM β-glycerol phosphate, 10mM Sodium orthovanadate, 10% Glyercol, 0.5mM EDTA, 0.5mM $MgCl_2$ , 500mM NaCl, 1mM DTT, 0.2% Triton-X, 1 tablet of EDTA-free mini cOmplete protease inhibitors (every 10mL lysis buffer), and 500units of Bensonase (every 1mL of lysis buffer). After the 100$\mu$L of lysis buffer is added to the cell pellet, the tube is then rotated at 4°C for 30min, then we sonicate for 5min (30sec on/30sec off). Lysates are quantified using BCA assay (Biorad). After quantification 30$\mu$g of lysate is fractionated on an SDS-PAGE gel, transferred, and blotted with primary, then secondary-HRP coupled antibodies. The RAG1, and RAG2 antibodies were kind gifts from the Schatz lab, and Tata binding protein was used as the loading control.

Halo-RAG1 and RAG2 *In Vivo* Pull-down.

To perform the Halo-RAG1 and RAG2 pull down experiment the Halo pull-down kit (Promega) was used. First an 80% confluent 10cm dish of 293T cells were transfected with 20$\mu$g of Halo-RAG1 or Halo-alone vector, and 20$\mu$g of RAG2 vector using BioT. The transfection proceeded for 48hrs then the cells were harvested. The cells were lysed using 50mM Tris (pH 7.5), 150mM NaCl, 1mM DTT and 1% NP-40 on ice for 30min, then centrifuged at 14,000 rpm for 2 mins *(29)*. The Halo-RAG1/RAG2 or Halo-alone/RAG2 lysates were added to Halo-link resin, and were rotated at 4°C for a 1hr and 30mins. The Halo-link resin was washed 3 times, and eluted using an SDS solution provided in the Halo pull-down kit. The eluate was then fractionated on an SDS-PAGE and blotted against RAG2.

2D single-molecule imaging, localization and tracking in live pro-B cells.

All 2D single-molecule imaging experiments were conducted using a Nikon Eclipse Ti microscope with a 100X oil-immersion objective lens (N.A. = 1.49) paired with NIS elements software. Highly-inclined and laminated optical (HILO) sheet imaging was performed for all live cell single-molecule experiments *(30)*. Experiments imaging Halo-RAG derivatives in the absence of the *IgH* locus used 0.3-10nM of JF549 dye depending on the expression level of Halo-RAG derivatives, which was excited with a 561 nm laser, and imaged with a 10ms or 20ms integration time. To image the *IgH* locus and Halo-RAG1 or Halo-RAG1/RAG2 simultaneously we recruited TET-R-GFP to Tet operator sites in the *IgH* locus, excited with a 488 nm laser, then we labeled Halo-RAG1 or Halo-RAG1/RAG2 with 75nM JF646 excited with a 647 nm laser and imaged continuously with a 10ms (for diffusion) or 20ms (for dwell-times) integration time. For single molecule tracking, the spot localization (x, y) was obtained through 2D Gaussian fitting based on the multiple-target tracing (MTT) algorithms using a home built Matlab program (SLIMfast) *(31)*. The tracking parameters are listed in the Table S1. For the analysis of RAG1 and RAG1/RAG2 diffusion, the maximum expected diffusion coefficient was set to $D = 3\mu m^2/s$. Diffusion coefficients were calculated for trajectories that lasted at least 5 consecutive frames, and only tracks with linear fits with an $R^2 \geq 0.8$ using the MSDanalyzer are included. When analyzing RAG1 and RAG1/RAG2 dwell-time data the maximum expected diffusion coefficient was set to $D = 0.05\mu m^2/s$ so we can investigate binding events *(11, 12)*. The dwell-time was directly calculated from the length of the trajectory, and only tracks that lasted at least 2 consecutive frames were included.

When comparing *IgH* locus and H2B dynamics we determined the extent to which apparent diffusion caused by localization uncertainty was confounding our measurements using homemade matlab code *(32)*. We found the apparent diffusion was smaller for *IgH* than for H2B, bolstering the observation of faster *IgH* diffusivity relative to H2B (Fig S10, A to D). For the *IgH* locus recruitment studies, as a first level of distinction RAG1 and RAG1/RAG2 trajectories within 3 pixels of the *IgH* locus are considered *IgH*-proximal, and were segregated from non-*IgH*-proximal trajectories using a matlab script (kind gift from Jens Schmidt) *(12)*. To measure long-lived RAG1 dwell-times genome-wide, and at the *IgH* locus two color imaging was performed using a 20ms integration time followed by 300ms darktime *(11, 12)*. To determine the long-lived RAG1 dwell-time at the *IgH* locus, the reslice tool in ImageJ was used on the channel for RAG1 (647nm)) and the channel for the *IgH* locus (488nm), and then merged to identify co-localization events. The long-lived dwell-time for RAG1 was directly calculated from the number of frames it remained bound. To determine the genome-wide long-lived RAG1 mean dwell-time, a single vertical reslice at the center of a pro-B cell spanning its diameter was taken and the trajectories present were measured, and this was repeated for multiple cells. Lastly, to determine the frequency, and probability of binding for RAG1 or RAG1/RAG2 to the *IgH* locus, we used the 10ms two-color imaging data. We used the reslice and measure tool in ImageJ on the channel for RAG1 or RAG1/RAG2 (647nm) and the channel for the *IgH* locus (488nm), and then merge to identify co-localization events. A total of n = 65 cells were assayed per experiment, and only one *IgH* allele was considered per cell. We determined the frequency of RAG1 or RAG1/RAG2 binding to the *IgH* locus by dividing the number of RAG1 or RAG1/RAG2 co-localization events at the *IgH* locus by the total number of *IgH* loci assayed. We determined the probability of binding for RAG1 and RAG1/RAG2 to the IgH locus by dividing the time RAG1 or RAG1/RAG2 spend bound to the *IgH* locus, by the total time the *IgH* locus spends bound and unbound (Total Time = 2000 frames/per cell). Three independent experiments were performed for RAG1 and RAG1/RAG2.

Supplemental Figure 1

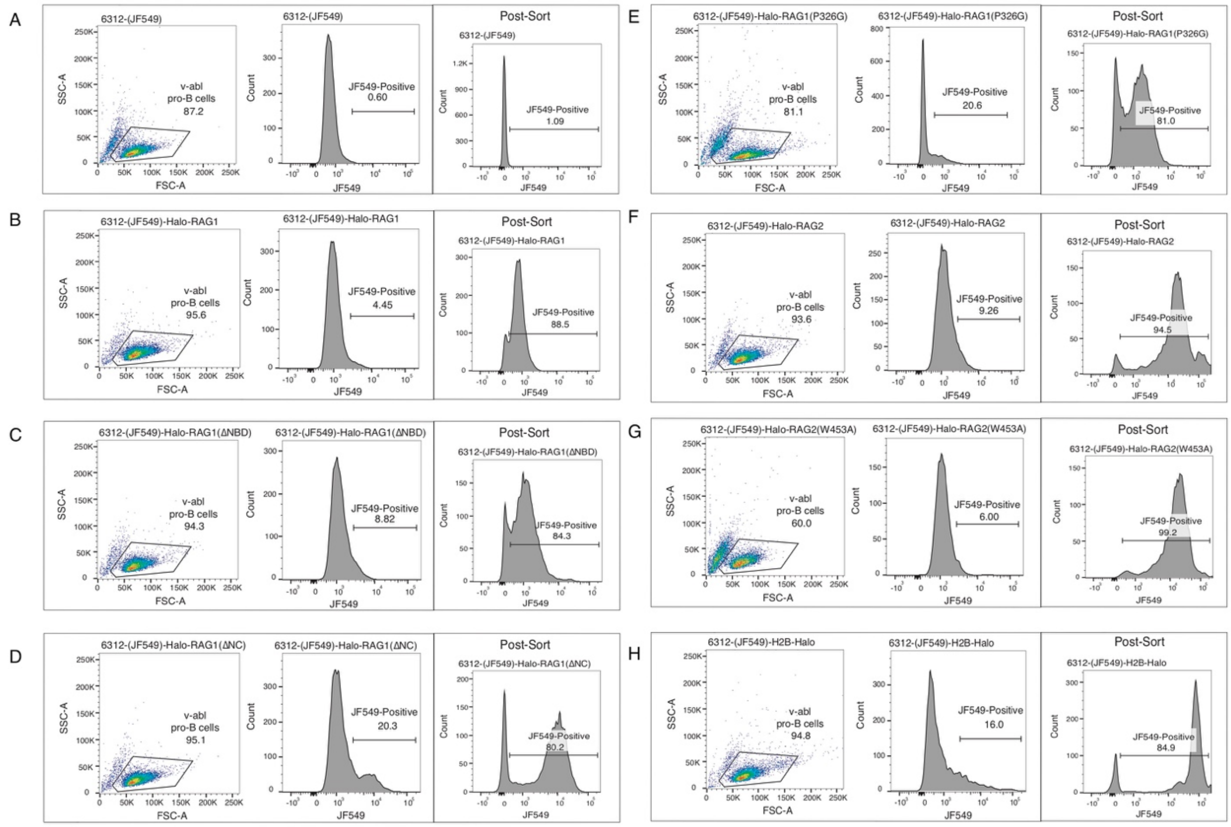


**Fig. S1. Retroviral infection efficiency of 6312 pro-B cells with Halo-RAG1, Halo-RAG1(ΔNBD), Halo-RAG1(ΔNC), Halo-RAG1(P326G), Halo-RAG2, Halo-RAG2(W453A) or H2B-Halo.** (**A**) FACs was performed on 6312 uninfected or (**B-H**)**.** infected with Halo-RAG1, Halo-RAG1(ΔNBD), Halo-RAG1(ΔNC), Halo-RAG1(P326G), Halo-RAG2, Halo-RAG2(W453A), and Halo-H2B. Cells to the right of the JF549 gate are positive for Halo. The infected cells were then sorted to purity, and the post sort histogram is shown in the third panel.

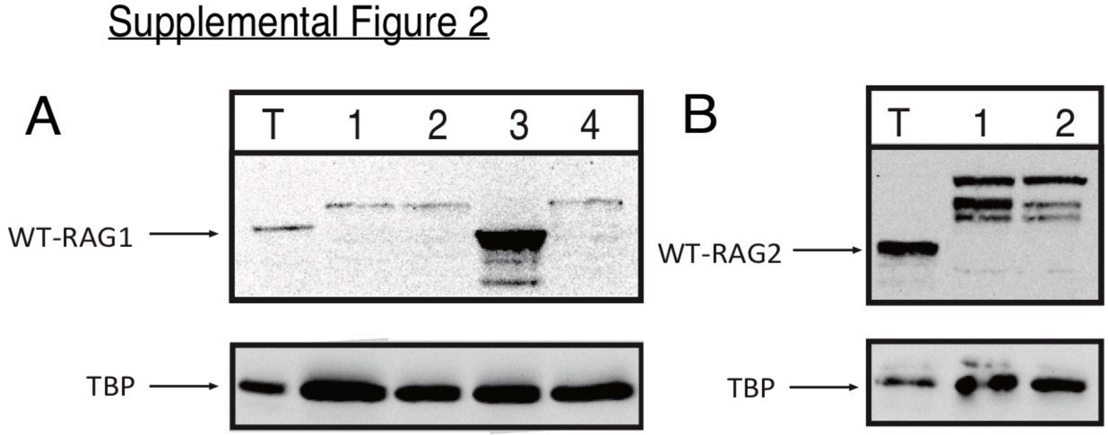


**Fig. S2. Western blot of Halo-RAG1, Halo-RAG1(ΔNBD), Halo-RAG1(ΔNC), Halo-RAG1(P326G), Halo-RAG2 or Halo-RAG2(W453A).** (**A**) We performed a RAG1 western blot after introducing the Halo-RAG1 mutants into 6312, and we found they were all intact (lane T: thymic lysate, lane 1: 6312-Halo-RAG1, lane 2: 6312-Halo-RAG1(ΔNBD), lane 3: 6312-Halo-RAG1(ΔNC), and lane 4: 6312-Halo-RAG1(P326G)). Tata-binding protein (TBP) was used as the loading control. (**B**) However once we introduced Halo-RAG2 mutants, and blotted against RAG2 we found they were degraded from their C-terminal ends (lane T: thymic lysate, lane 1: 6312-Halo-RAG2, and lane 2: 6312-Halo-RAG2(W453A)). TBP was used at the loading control.

#### Supplemental Figure 3

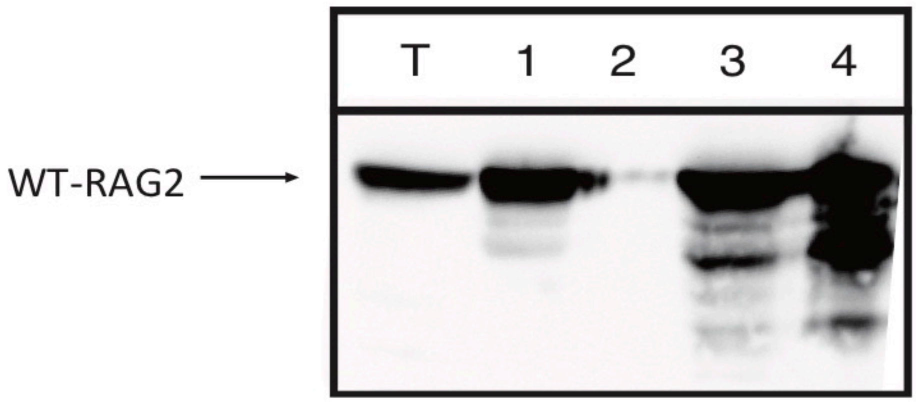


**Fig. S3.** ***In vivo*** **pull-down assay of RAG2 by Halo-RAG1.** To confirm that Halo-RAG1 can interact with RAG2, we performed an *in vivo* Halo pull-down experiment, and then performed a western blot against RAG2, using a RAG2-antibody. We found Halo-RAG1 (lanes 3 and 4) can indeed interact with RAG2, but Halo alone (lanes 1 and 2) does not (lane T: thymic lysate, lane 1: Halo/RAG2 input, lane 2: Halo/RAG2 pull-down, lane 3: Halo-RAG1/RAG2 input, and lane 4: Halo-RAG1/RAG2 pull-down).

Supplemental Figure 4

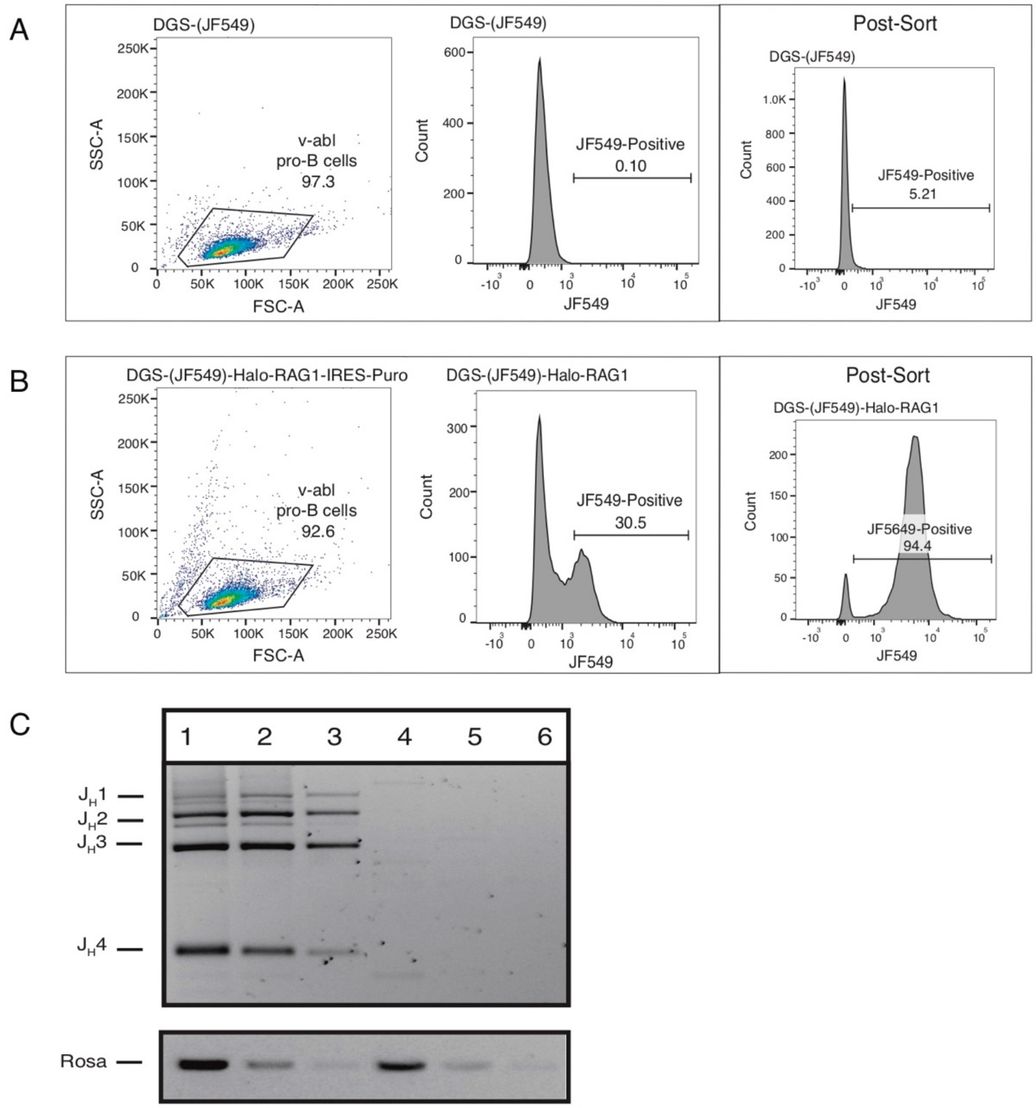


**Fig. S4. Retroviral infection efficiency of DGS pro-B cells with Halo-RAG1-IRES-Puromycin after selection, and DJ recombination.** In order to introduce Halo-RAG1 into DGS we created a Halo-RAG1-IRES-Puromycin vector, because the Halo-RAG1 alone vector was silenced for reasons that remain unclear. (**A**) FACs of DGS uninfected, and (**B**) infected cells (after puromycin selection). Cells to the right of the JF549 gate are positive for Halo. The infected cells were then sorted to purity, and the post-sort histogram is shown in the third panel of A and B. After introducing Halo-RAG1 into DGS, a RAG1-deficient pro-B cell line, we performed a DJ recombination assay to confirm Halo-RAG1 is functional in recombination. We treated DGS-Halo-RAG1 or DGS alone with STI-571 for 48hrs. (**C**) We find that we only observe DJ recombination products once Halo-RAG1 is present. The *Rosa* locus was used as the loading control. (lane 1: DGS-Halo-RAG1-48hr STI, lane 2: 1/5 dilution of DGS-Halo-RAG1-48hr STI, lane 3: 1/25 dilution of DGS-Halo-RAG1-48hr STI, lane 4: DGS-48hr STI, lane 5: 1/5 dilution of DGS-48hr STI, and lane 6: 1/25 dilution of DGS-48hr STI).

Supplemental Figure 5

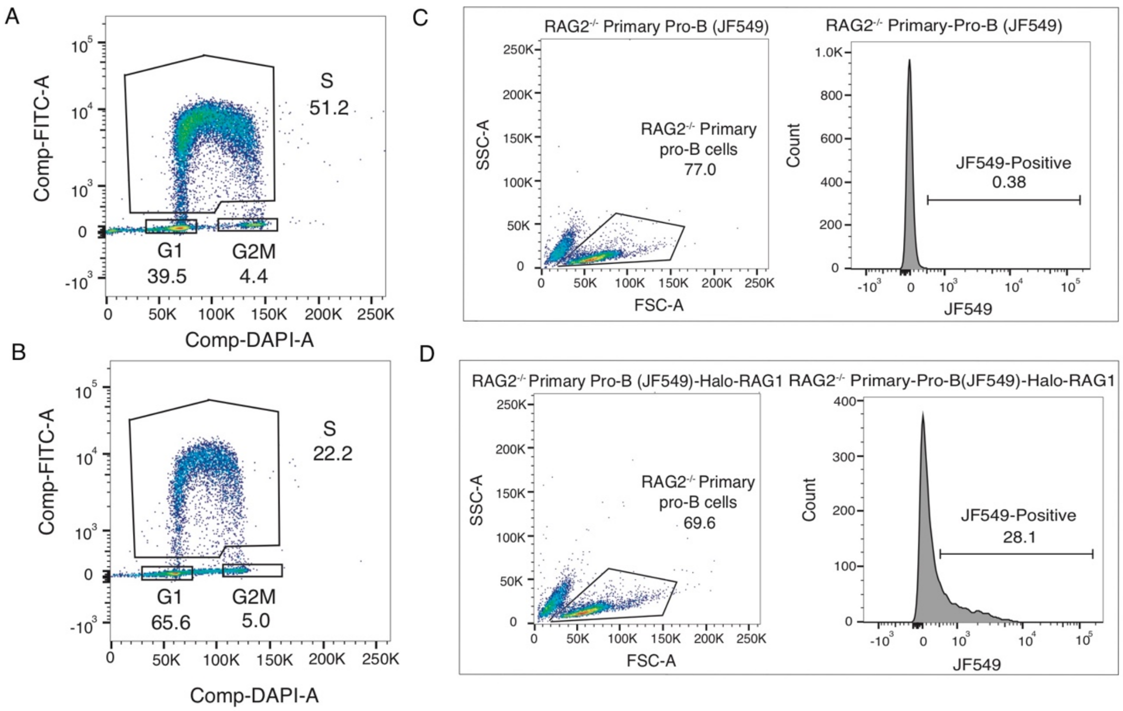


**Fig. S5. Effect of STI treatment on 6312-Halo-RAG1 cell cycle distribution, and retroviral infection of RAG2-deficient primary pro-B cells with Halo-RAG1.** To determine the distribution of 6312-Halo-RAG1 cells in the G1, S and G2/M phases of the cell cycle (**A**) without or (**B**) with STI-571 treatment we labeled the cells with EdU, then DAPI and performed a FACs assay. EdU is incorporated into the genome by cells in the S phase of the cell cycle, and FITC labeled picolyl azide is added, which covalently couples to EdU. The y-axis on the FACs plots represent EdU, and the x-axis represents DAPI. (top gate: S phase, bottom-left: G1 phase, and bottom right: G2/M phase). Next to introduce Halo-RAG1 in RAG2-deficient primary pro-B cells, we performed a retroviral transduction, and then added JF549 to label Halo-RAG1, and performed FACs to determine the infection efficiency. (**C**) Uninfected and (**D**) infected RAG2-deficient FACs plots are shown. The fraction of cells to the right of the JF549 gate are halo positive.

#### Supplemental Figure 6

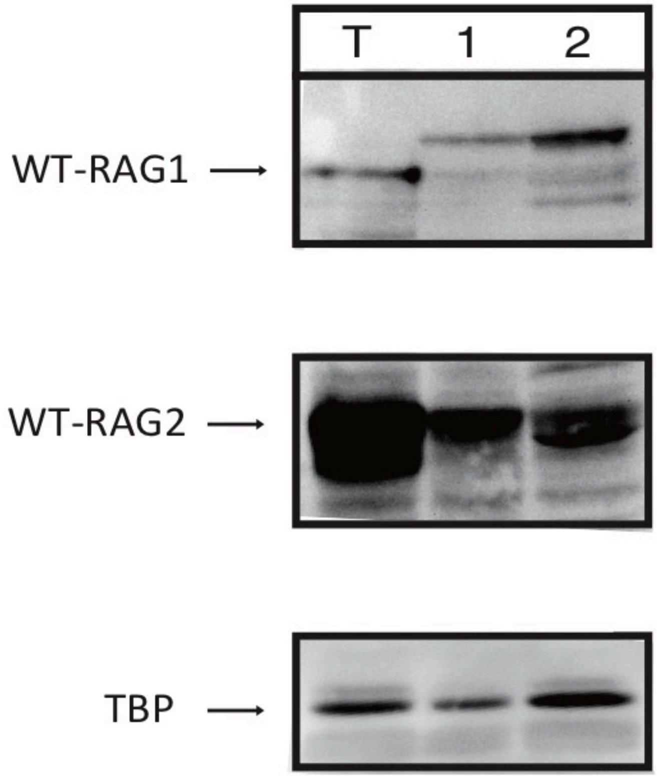


**Fig. S6. Western blots of 6312-Halo-RAG1 + RAG2 or RAG2(W453A).** To introduce unlabeled RAG2 and RAG2(W453A) in the presence of Halo-RAG1 we performed a lentiviral transduction, followed by puromycin selection. We then performed western blots to confirm RAG2 and RAG2(W453A) were introduced into 6312-Halo-RAG1 (lane T: thymic lysate, lane 1: 6312-Halo-RAG1/RAG2, and lane 2: 6312-Halo-RAG1/RAG2(W453A)). We blotted against RAG1 (upper panel), RAG2 (middle panel), and TBP (lower panel).

### Supplemental Figure 7

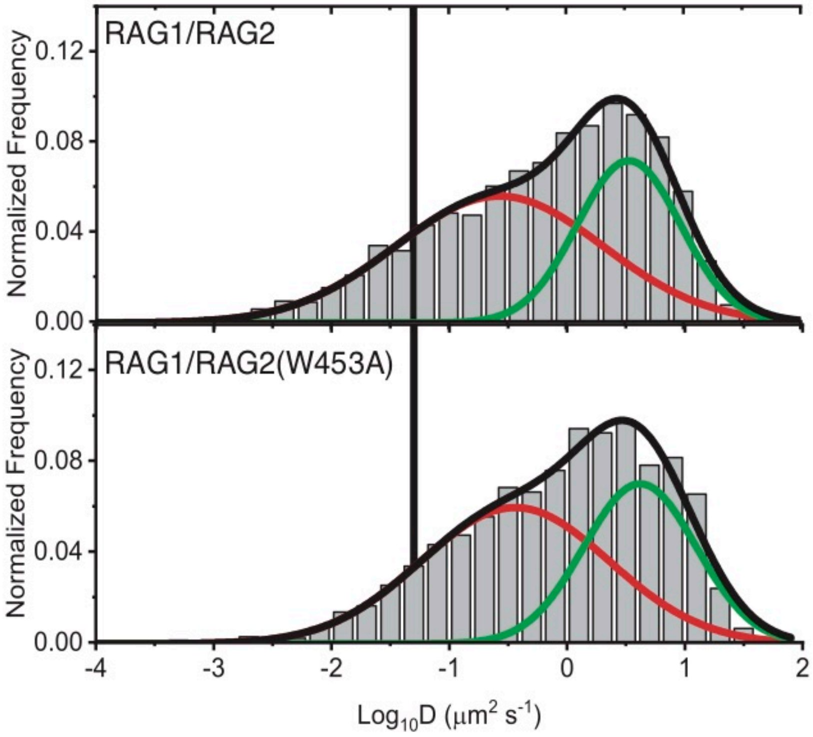


**Fig. S7. Representative diffusion coefficient distribution and Gaussian fits for 6312-Halo-RAG1/RAG2 and 6312-Halo-RAG1/RAG2(W453A).** Diffusion coefficient distributions and Gaussian fits for 6312-Halo-RAG1/RAG2 and 6312-Halo-RAG1/RAG2(W453A) (slow diffusion: red, and fast diffusion: green). The fraction of the histogram to the left of the black vertical line is slow diffusing based on the histone H2B slow diffusion coefficient cutoff.

#### Supplemental Figure 8

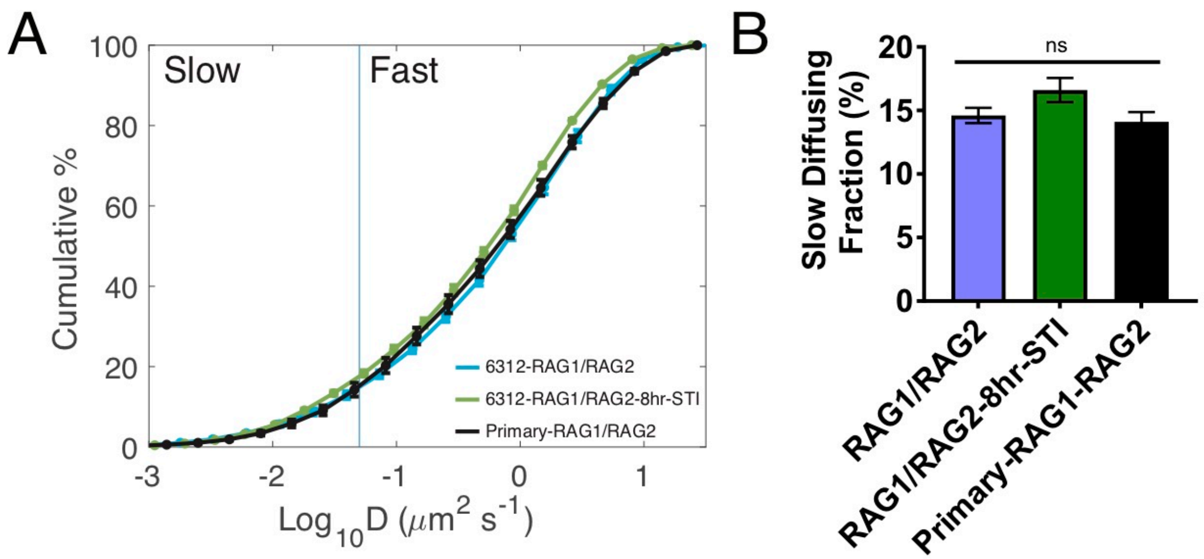


**Fig. S8. CDFs, and bound fraction of 6312-Halo-RAG1/RAG2, 6312-Halo-RAG1/RAG2-8hr STI treatment, and RAG1-deficient primary-Halo-RAG1/RAG2.** (**A**)**.** CDFs of 6312-Halo-RAG1/RAG2 (cyan), 6312-Halo-RAG1/RAG2-STI-8hr (green), and RAG1-deficient-Halo-RAG1/RAG2 (24hr IL-7 withdrawal) (black). (**B**) Bound fraction of 6312-Halo-RAG1/RAG2, 6312-Halo-RAG1/RAG2-STI-8hr, and RAG1-deficient-Halo-RAG1/RAG2 (24hr IL-7 withdrawal). The fraction of the CDF to left of the blue vertical line is bound. The error bars represent the SEM.

Supplemental Figure 9

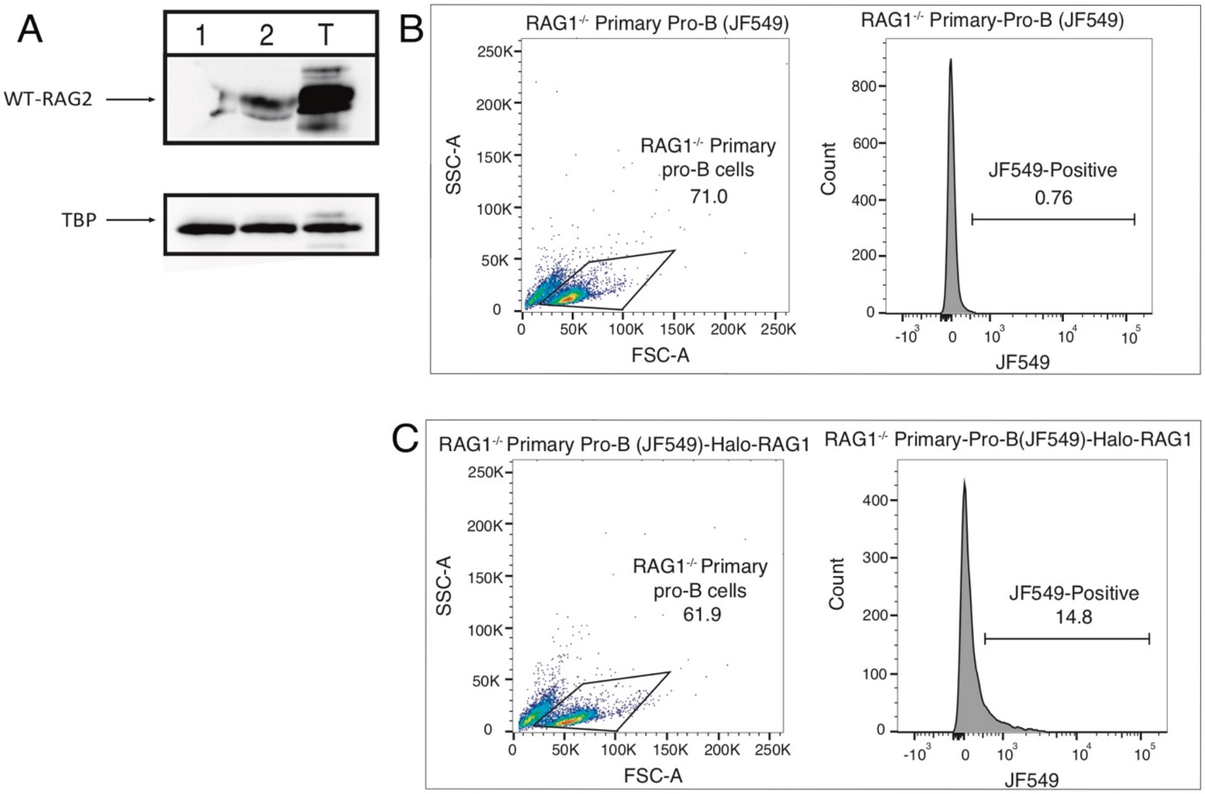


**Fig. S9. Retroviral Infection efficiency of RAG1-deficient primary pro-B cells with Halo-RAG1, and western blot of RAG2 induction.** (**A**) Western blot of RAG2 induction in RAG1-deficient primary pro-B cells (lane 1: IL-7 present, lane 2: 24hr IL-7 withdrawal, and lane T: Thymic lysate). (**B**) Uninfected RAG1-deficient primary pro-B cells. (**C**) Infected RAG1-deficient primary pro-B cells with Halo-RAG1. Cells to the right of the JF549 gate are positive for Halo.

Supplemental Figure 10

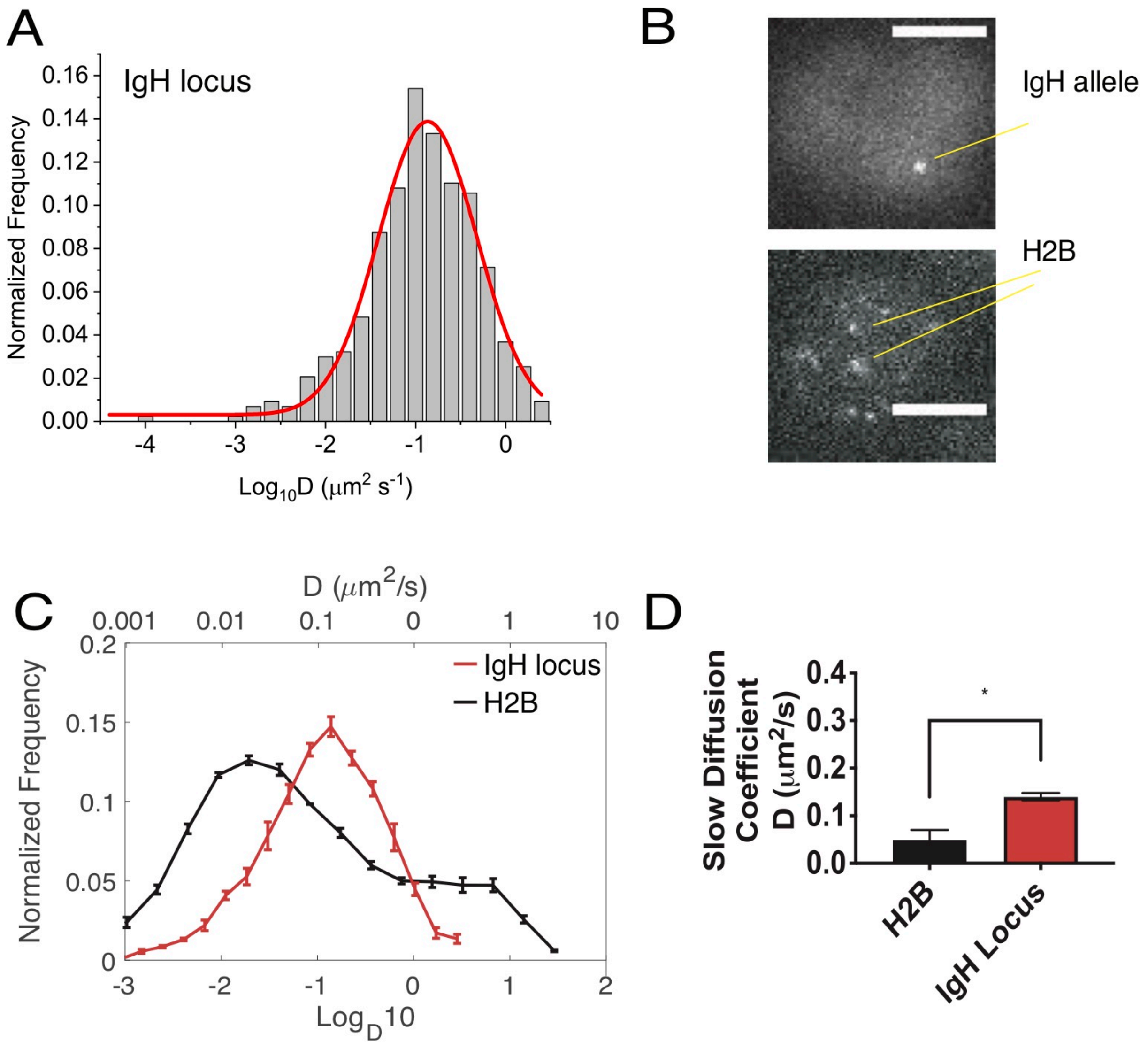


**Fig. S10. Comparison of H2B and *IgH* locus diffusivity.** (**A**) Representative diffusion coefficient distribution for the *IgH* locus with Gaussian fit (red). (**B**) Images of IgH and H2B (scale bar = 5μm). (**C**) Diffusion coefficient histogram overlay for H2B (black) and the *IgH* locus (red). (**D**) Mean slow diffusion coefficient for H2B (black) and the *IgH* locus (red). (Error bars represent SEM, *p < 0.05).

Supplemental Figure 11

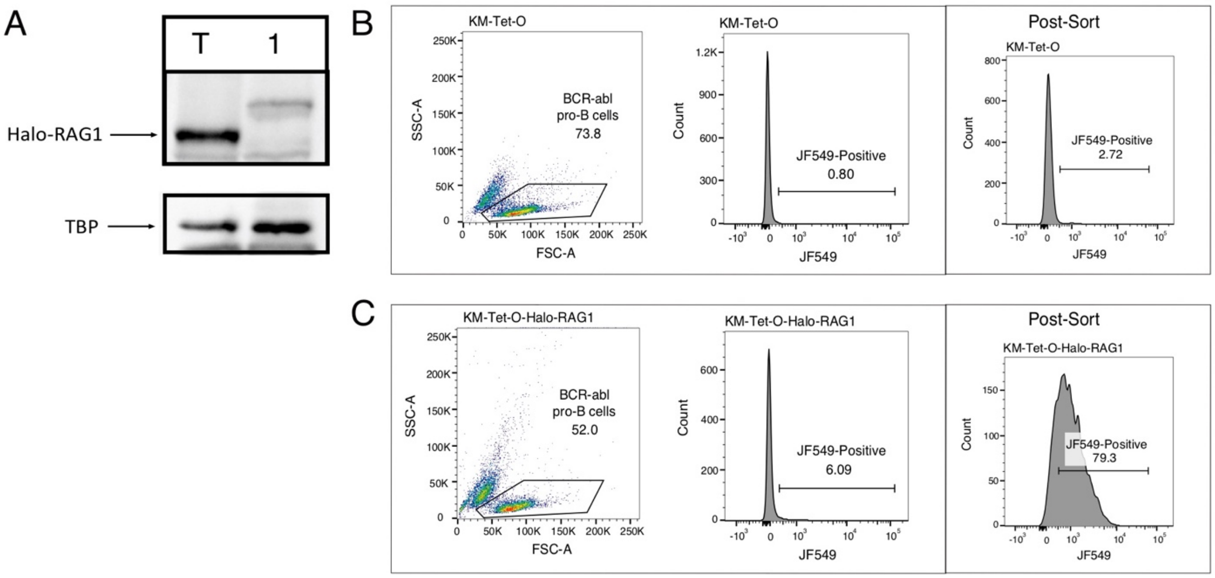


**Fig. S11. Retroviral infection efficiency and western blots of KM-Tet-O pro-B cells with Halo-RAG1.** (**A**) RAG1 western blot of Halo-RAG1 in the KM-Tet-O pro-B cell line. (**B**) Uninfected KM-Tet-O pro-B cells, and (**C**) Infected KM-Tet-O pro-B cells with Halo-RAG1. Cells to the right of the JF549 gate are positive for Halo. The infected cells were then sorted to purity, and the post-sort histogram is shown in the third panel of B and C.

Supplemental Figure 12

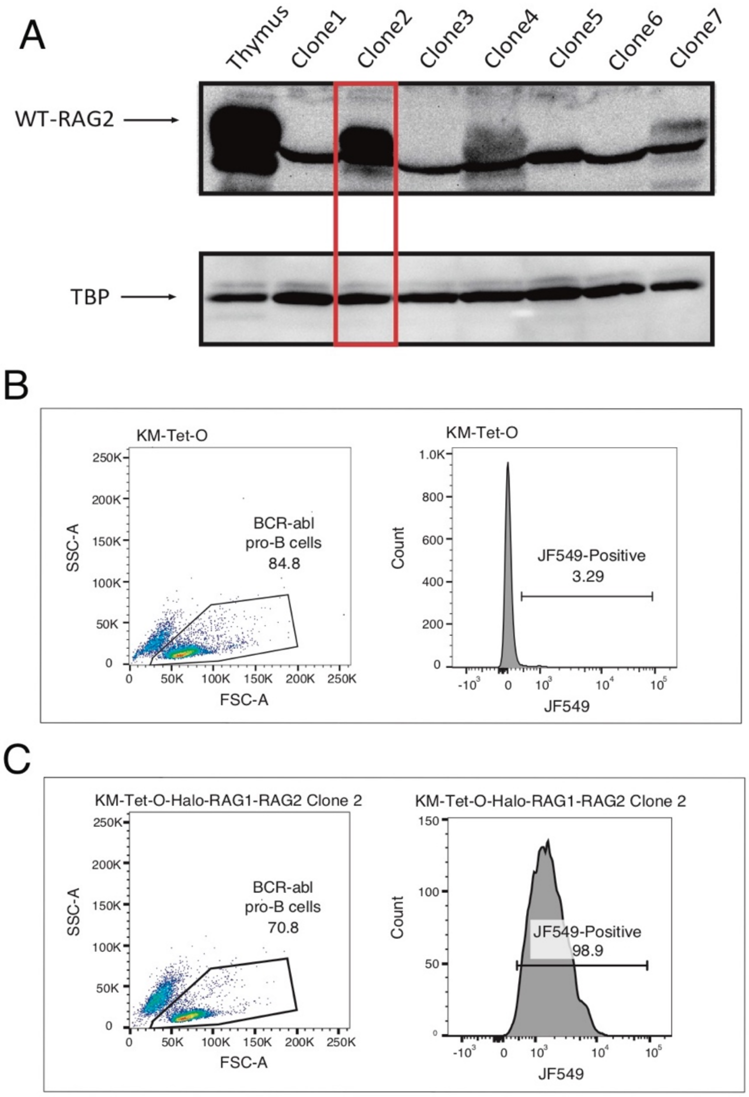


**Fig. S12. Western blots, and FACs of KM-Tet-O-Halo-RAG1/RAG2 in pro-B cells.** To introduce RAG2 into KM-Tet-O-Halo-RAG1 we performed lentiviral transduction using RAG2-IRES-Puromycin. Puromycin selection was performed however, the puromycin concentration required for growth did not cause significant cell death, therefore a large fraction of uninfected cells persisted. (**A**) We then single cell cloned the infected cells, blotted against RAG2, and identified a high expressing clone (lane 2, red box). We then performed FACs on the cell line to confirm the clone also harbored Halo-RAG1. (**B**) FACs plots of uninfected KM-Tet-O-pro-B cells, and (**C**) infected KM-Tet-O-Halo-RAG1/RAG2 pro-B cells. Cells to the right of the JF549 gate are positive for Halo.

#### Supplemental Figure 13

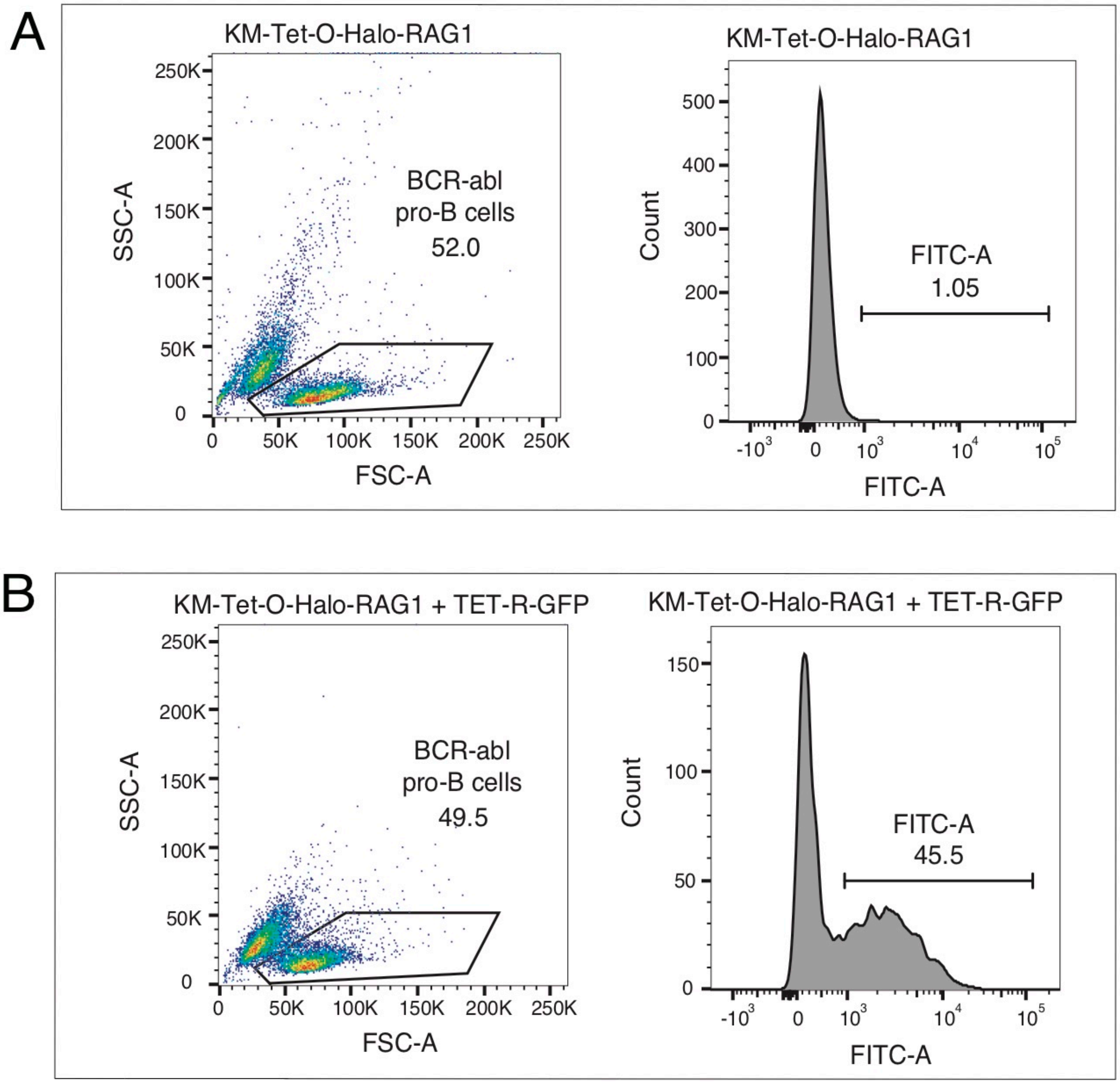


**Fig. S13. Retroviral infection efficiency of KM-Tet-O-Halo-RAG1 pro-B cells with Tet-R-GFP.** (**A**) FACs of uninfected KM-Tet-O-Halo-RAG1 pro-B cells, and (**B**) TET-R-GFP infected KM-Tet-O-Halo-RAG1 pro-B cells. Cells to the right of the FITC gate are positive for TET-R-GFP.

<u>Supplemental Figure 14</u>

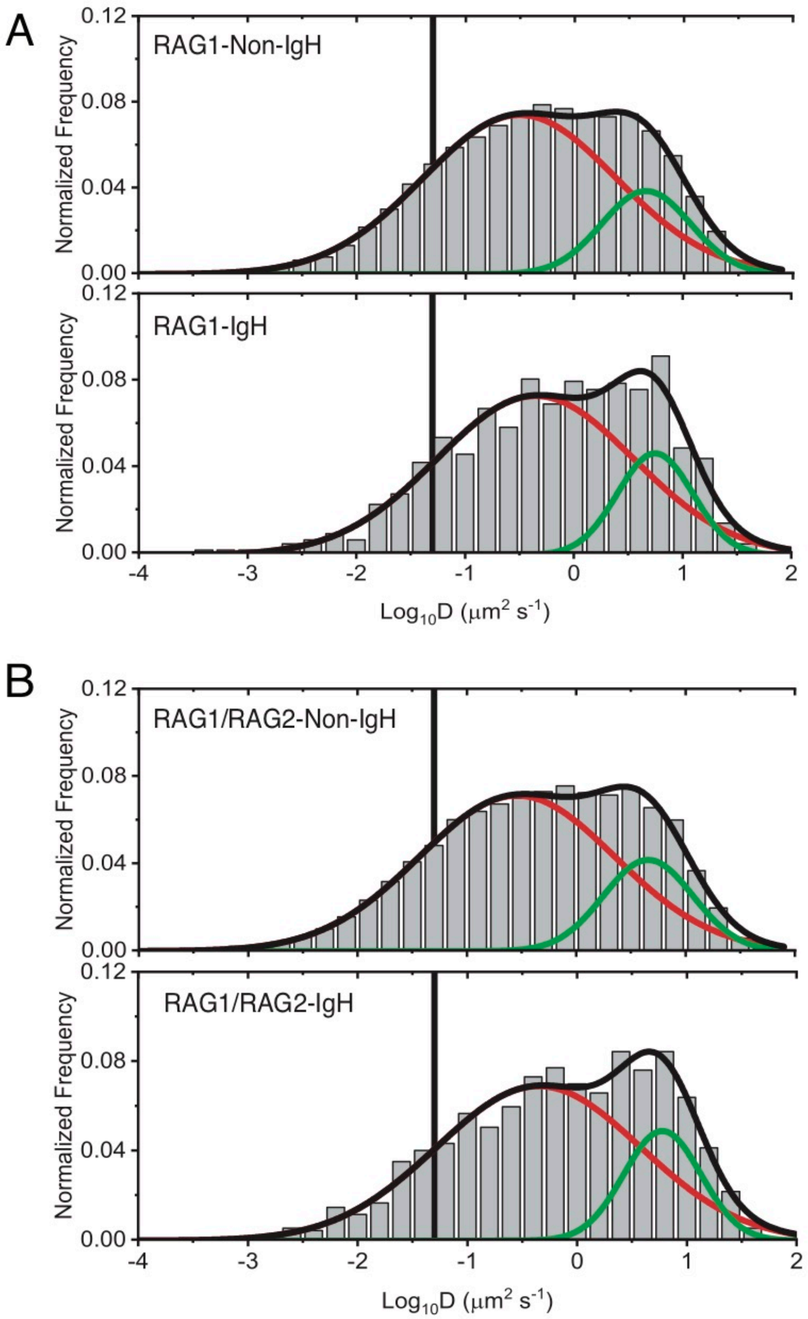


**Fig. S14. Gaussian Fits to *IgH* and Non-*IgH*-associated Halo-RAG1 or Halo-RAG1/RAG2 diffusion coefficient distributions.** (**A**) Gaussian fits of *IgH*-proximal, and non-*IgH*-proximal Halo-RAG1 trajectories. (**B**) Gaussian fits of *IgH*-proximal, and non-*IgH*-proximal Halo-RAG1/RAG2 trajectories. The fraction of the histogram to the left of the black vertical line is slow diffusing based on the histone H2B slow diffusion coefficient cutoff.

#### Supplemental Figure 15

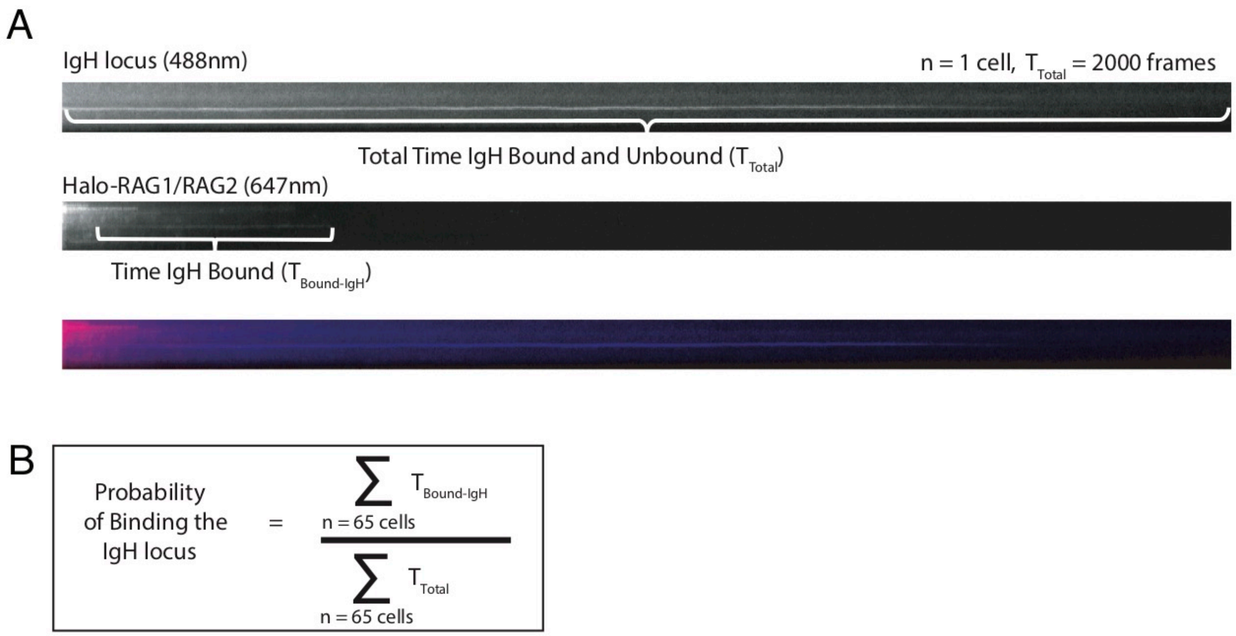


**Fig. S15. Computing the probability of binding to the *IgH* locus** (**A**) Resliced kymographs of 10ms two-color imaging data of the IgH locus (top), Halo-RAG1/RAG2 (middle), and merge (bottom) recorded for 2,000 frames. If the *IgH* locus diffused out of the field of view during the length of the recording it was defined as unbound. (**B**) Equation used to compute the probability of binding to the *IgH* locus. Since $T_{Total}$ = 2,000 frames per cell, the only parameter varying in the equation is $T_{Bound\text{-}IgH}$. Three independent experiments are performed, and 65 cells are assayed per experiment.

**Table S1. Parameters for 2D single-molecule imaging, tracking and analysis**

| Parameter | Diffusion Analysis | Dwell-time Analysis |
|---|---|---|
| Exposure time (ms) | 10 | 20 |
| Excitation (nm) | 561 or 647 | 561 or 647 |
| Emission (nm) | 590 or 664 | 590 or 664 |
| Pixel size (nm) | 160 | 160 |
| Numerical Aperture | 1.49 | 1.49 |
| Expected $D_{max}$ ($\mu m^2/s$) | 3 | 0.05 |

**Movie S1. 10ms Tracking of H2B-Halo in live 6312-pro-B cells.** 2D tracking of H2B-Halo in the v-Abl pro-B cell line 6312. Images were collected using a 10ms integration time, and the cell permeable JF549 dye was used for visualization. The movie is time-stamped in milliseconds. The frame rate is set to 20 frames/sec for clarity.

**Movie S2. 10ms Tracking of Halo-RAG1 in live 6312-pro-B cells.** 2D tracking of Halo-RAG1 in the v-Abl pro-B cell line 6312. Images were collected as described in Movie S1. The movie is time-stamped in milliseconds. The frame rate is set to 20 frames/sec for clarity.

**Movie S3. 10ms Tracking of Halo-RAG1 in RAG2-deficient primary pro-B cells.** 2D tracking of Halo-RAG1 in RAG2-deficient primary pro-B cells. Images were collected as described in Movie S1. The movie is time-stamped in milliseconds. The frame rate is set to 20 frames/sec for clarity.

**Movie S4. 20ms Tracking of Halo-RAG1 in live 6312-pro-B cells.** 2D tracking of Halo-RAG1 in the v-Abl pro-B cell line 6312. Images were collected using a 20ms integration time, and the cell permeable JF549 dye was used for visualization The movie is time-stamped in milliseconds. The frame rate is set to 20 frames/sec for clarity.

**Movie S5. 10ms Tracking of Halo-RAG1/RAG2 in live 6312-pro-B cells.** 2D tracking of Halo-RAG1/RAG2 in the v-Abl pro-B cell line 6312. Images were collected as described in Movie S1. The movie is time-stamped in milliseconds. The frame rate is set to 20 frames/sec for clarity.

**Movie S6. 10ms Tracking of Halo-RAG1/RAG2 in live RAG1-deficient primary pro-B cells (24hr IL-7 Withdrawal).** 2D tracking of Halo-RAG1/RAG2 in RAG1-deficient primary pro-B cells. Images were collected as described in Movie S1. The movie is time-stamped in milliseconds. The frame rate is set to 20 frames/sec for clarity.

**Movie S7. 20ms Tracking of Halo-RAG1/RAG2 in live 6312-pro-B cells.** 2D tracking of Halo-RAG1/RAG2 in the v-Abl pro-B cell line 6312 treated for 8hr with STI-571. Images were collected as described in Movie S8. The movie is time-stamped in milliseconds. The frame rate is set to 20 frames/sec for clarity.

**Movie S8. 10ms Tracking of Halo-RAG1/RAG2 and the *IgH* locus in live KM-Tet-O-pro-B cells.** 2D tracking of Halo-RAG1/RAG2 and the *IgH* locus in the bcr-Abl pro-B cell line KM-Tet-O. Images were collected using a 10ms integration time, the JF646 dye was used to illuminate Halo-RAG1/RAG2, and TET-R-GFP was used to visualize the *IgH* locus. The frame rate is set to 30 frames/sec for clarity.

**Movie S9. 20ms Tracking of Halo-RAG1/RAG2 and the *IgH* locus in live KM-Tet-O-pro-B cells.** 2D tracking of Halo-RAG1/RAG2 and the *IgH* locus in the bcr-Abl pro-B cell line KM-Tet-O. Images were collected using a 20ms integration time, the JF646 dye was used to illuminate Halo-RAG1/RAG2, and TET-R-GFP was used to visualize the *IgH* locus. The frame rate is set to 30 frames/sec for clarity.

**Movie S10. 20ms integration time and 300ms dark time Tracking of Halo-RAG1 and the *IgH* locus in live KM-Tet-O-pro-B cells.** 2D tracking of Halo-RAG1 and the *IgH* locus in the bcr-Abl pro-B cell line KM-Tet-O. Images were collected using a 20ms integration time/ 300ms dark time, the JF646 dye was used to illuminate Halo-RAG1, and TET-R-GFP was used to visualize the *IgH* locus. The frame rate is set to 15 frame/sec for clarity.
